# Scale-dependent tipping points of bacterial colonization resistance

**DOI:** 10.1101/2021.05.13.444017

**Authors:** Yuya Karita, David T. Limmer, Oskar Hallatschek

## Abstract

Bacteria are efficient colonizers of a wide range of secluded micro-habitats, such as soil pores, skin follicles, or intestinal crypts. How the structural diversity of these habitats modulates microbial self-organization remains poorly understood, in part because of the challenge to specifically manipulate the physical structure of microbial environments. Using a microfluidic device to grow bacteria in crypt-like incubation chambers of systematically varied lengths, we show that small variations in the physical structure of the micro-habitat can drastically alter bacterial colonization success and resistance against invaders. Small crypts are un-colonizable, intermediately sized crypts can stably support dilute populations, while beyond a second critical lengthscale, populations phase-separate into a dilute and a jammed region. The jammed state is characterized by extreme colonization resistance, even if the resident strain is suppressed by an antibiotic. Combined with a flexible biophysical model, we demonstrate that colonization resistance and associated priority effects can be explained by a crowding-induced phase transition, which results from a competition between proliferation and density-dependent cell leakage. The emerging sensitivity to scale underscores the need to control for scale in microbial ecology experiments. Systematic flow-adjustable lengthscale variations may serve as a promising strategy to elucidate further scale-sensitive tipping points and to rationally modulate the stability and resilience of microbial colonizers.

---

Natural microbial communities are often found to be remarkably stable, capable of either quickly recovering from disturbances or remaining essentially unaffected by them^1–4^. Stability is particularly puzzling in small populations, which are prone to number fluctuations and lack the size and extent to buffer against local environmental changes. Nevertheless, small but stable populations have been found in association with spatially defined micro-habitats^4–10^.

Strains that colonize cavities are sometimes found to be so stable that they hold their ground against even much fitter invaders^11^. For example, *Bacteroides fragilis* is a particularly resilient colonizer of crypts in mouse guts^7^. Conspecifics are unable to invade, unless the resident strain is strongly suppressed by an antibiotic. A similar colonization resistance has been demonstrated for groups of ceca microbiota in mice guts^9^ and for *Lactobacillus plantarum* in fly guts^10, 12^.

The ubiquity of micro-habitat associated stability and colonization resistance raises the question of whether these features generically emerge in confined spaces, for example soil pores^13–15^, skin follicles^4, 16^, or crypts and folds in gut-like environments^5,17,18^. Previous studies have identified biological features, such as suppressed biofilm growth or the expression of specific adhesion molecules, that promote stability in specific systemsb^1, 7, 19–21^. However, we currently lack systematic scale-dependent measurements to identify a generic mechanism of stability and resilience in micro-habitats, as well as a theory that could predict colonization success and tipping points. To fill this gap, we developed an approach to measure the scale-dependence of microbial colonization patterns combined with a predictive theory of how microbes invade, occupy and protect confined micro-habitats.

## Experimental Setup

Our experiments employ a microfluidic incubation device that allows to continuously monitor bacterial population dynamics in crypt-shaped chambers across many lengthscales (Fig. 1a). A supply channel is used to continuously perfuse the device with media enabling the experiments to run under constant conditions for several days. As bacteria are inoculated and pass through the supply channel, they get exposed to rectangular cavities of systematically varied depths (10-350 μm). Even though the fluid inside these cavities is largely stagnant, it is nutrient rich and hence supports growth, due to the rapid diffusion of small nutrient molecules from the supply channel^22, 23^.

**Figure 1.**
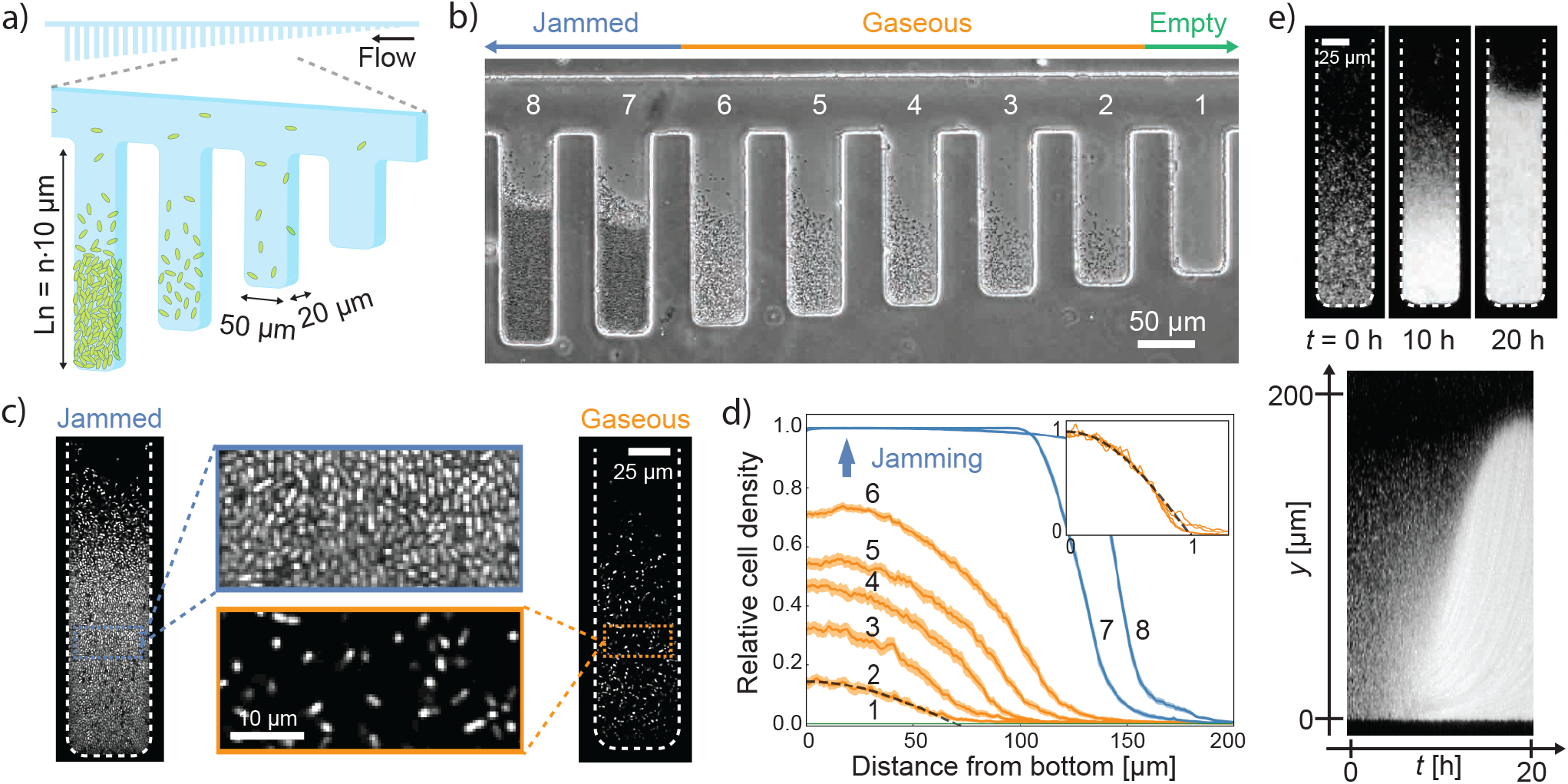
Microfluidic experiments reveal lengthscale-dependent colonization patterns. (a) A scheme of our Microfluidic Panflute incubation device: Rectangular cavities of systematically varied depths (*n* = 1, 2,…, 35) are connected to a common supply channel through which media and bacteria flow. (b) The steady state after five days of incubation of a fly gut bacterium (*A. indonesiensis*). Depending on their length, cavities could not be invaded (1), hosted a gaseous population (2-6) or a phase-separated population with a jammed and gaseous state (7-8). (c) Confocal images of a partially jammed and gaseous population. The zoomed-in images are magnifications of the zoomed-out snapshots. (d) Steady-state cell density profiles obtained from time lapse movies. The shaded regions show the standard error of the mean. The profiles of gaseous phases (orange) collapsed to our linearized establishment model (black) upon rescaling both axes (inset). (e) A kymograph of the jamming front movement.

In this device, lengthscale-dependent ecological processes can be identified by comparing the colonization dynamics across the sequence of chambers. To capture the differential population dynamics in single microscopy frames, we ordered the cavities according to size (see Fig. S1 for a randomized control). The device thus resembles a panflute in appearance, so we refer to our device as a “Microfluidic Panflute”. We employed it to explore the colonization dynamics of several bacterial genera, focusing mainly on *Acetobacter*, which is prevalent in the fly gut^10, 24^ and grows aerobically.

## Results

We found that the emerging population dynamics sensitively depend on the length of the incubation chamber (Fig. 1b and Supplementary Movie 1). The scale sensitivity is particularly strong near two recognizable phase transitions:

### Establishment Transition

While all cavities are sporadically visited by cells, colonization attempts remain unsuccessful in small chambers. In chambers exceeding a certain threshold length (170 μm in Fig. 1b), cell densities stabilize after 2-3 days of incubation and are maintained for at least five days. Cell densities, as measured from the time-averaged signal intensity, increase with chamber length, are highest at the floor of the cavities and gradually decay towards a line of zero density (Fig. 1d). We call this regime “gaseous” because the cell packing fraction is small and cells diffuse almost freely (SI Fig. S2).

### Jamming transition

When the chamber length exceeds a second threshold (220 μm in Fig. 1b), a densely populated region appears at the bottom of the cavities that is sharply separated from a gaseous region towards the opening of the cavities (see chambers 6 and 7 in Fig. 1b). Confocal imaging shows that neighboring cells are in direct contact in the dense phase, which is why we call the condensed phase “jammed” (Fig. 1c). Dynamically, the jammed phase grows like a wave from the floor towards the open boundary of a chamber, as can be seen in the kymograph Fig. 1e. The growth of this wave slows down near the jamming transition (Supplementary Movie S1). Interestingly, the transition from gaseous to jammed is abrupt in the size of the chambers. Between two neighboring cavities, differing by just 5% in length, the colonization state transitions from gaseous to nearly 75% jammed (quantified in Fig. 1d).

We observed qualitatively similar colonization patterns for species of other genera, including *V. cholerae* and *L. lactis* (Fig. S3). We therefore sought to explain the pronounced lengthscale-sensitivity by a general species-independent mechanism.

### Linear Establishment Model

The colonization of a cavity can be viewed as a tug of war between cell proliferation and cell removal by outflow or death^1^. This competition can be considered in the absence of regulation or specific cell-cell interactions, in order to discern whether the rich scale-dependent phase behavior seen in our experiments is a consequence of general biophysical processes. To describe how the cell density *c*(*y*,*t*) at a vertical position *y* and time *t* changes over time, we use the linear reaction-diffusion equation 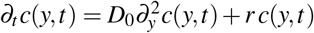 where the first term represents cell diffusion with diffusivity *D*_0_ and the second term represents cell proliferation with growth rate *r*. Since cells cannot penetrate the floor of the chamber, we use a reflecting boundary condition at *y* = 0, *∂_y_c*(0, *t*) =0. We also introduce an absorbing boundary at *y* = *L* where the cells are swept away by the media flow, *c*(*L*, *t*) = 0.

Our mathematical analysis (SI Sec. A.3) shows that the dynamics of the density profile can be decomposed into a sum of independently evolving normal modes. The empty state is stable if the amplitude of all normal modes shrink, which requires that the scale *L* of the population does not exceed the critical scale 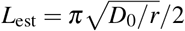. In turn, this implies that bacteria can establish in a chamber only if *L* > *L*_est_. Thus, establishment is promoted by increasing the growth rate or decreasing the diffusivity, which drives the cell leakage. Using the measured growth rate, *r* ≈ 0.33 ±0.01 h^−1^ (Fig. S4a and d), and diffusivity, *D*_0_ ≈ (0.37 ±0.01) × 10^3^ μm^2^/h (Fig. S2), we estimate establishment in our experiments to occur at the scale *L*_est_ ≈ 53 ± 1 μm. This is consistent with the empirical value 53 ±7 μm that we extrapolate from our measurements (Fig. S5b). We also confirmed that the establishment length changes predictably with variations in growth rate (Fig. S6). More importantly, the measured density profiles agree well with the cosine shape of the first normal mode, as observed in Fig. 1d, which is expected to dominate close to the onset of colonization (SI Sec. A.4). Our analysis is best suited to describe the bulk of the population where cell motion is dominated by diffusion. Deviations are expected, and indeed visible around the opening of cavities (near vanishing cell density) where the flow of the media is not negligible.

### Nonlinear Population Control

Our linear model can tell us whether bacteria grow in empty chambers but it remains blind to how a population of successful colonizers reaches a steady state with a finite population size and how stable this state is. To predict the long-term dynamics, we needed to include a (non-linear) population control term that modulates the competition between cell proliferation and removal. For example, bacterial batch cultures are often limited by nutrient deprivation or waste product accumulation, implying that the growth rate is not constant but decays with density (logistic population control). However, growth rates in the jammed and dilute phase were statistically indistinguishable (Fig. S4), suggesting that nutrient deprivation did not limit population growth. Therefore, we hypothesized that, while the growth rate remains approximately constant, the population outflow adjusts itself via a density-dependent diffusivity *D*(*c*). Steady state is reached when the cell leakage matches the influx of newborn cells in the bulk of the chamber.

### Crowding-induced Phase Transition

Our mathematical analysis shows that a monotonically increasing *D*(*c*) (more cells →more outflow) is capable of stabilizing a gaseous state inside the chambers (SI Sec. A.4). However, to reproduce a sudden jamming transition, *D*(*c*) has to have a region of negative slope at high densities (more cells →less outflow). Intuitively, this generates a positive feedback cycle. As the density fluctuates up, diffusion-induced outflow goes down, which leads to even higher cell densities, suppressing outflow even more and so on. The cycle only breaks when the bacteria jam and come into contact, upon which the bulk modulus and, hence, *D*(*c*) shoot up by several orders of magnitude^25^.

The required negative slope of *D*(*c*) could be induced at high density by constitutive or crowding-induced stickiness between cells, or active motility, which has been shown to drive phase-separation^26^. Our simulations (Fig. 2) and analytical arguments (SI Sec. A.7) show that even purely repulsive non-motile spheres exhibit a qualitatively similar phase behavior as seen in our experiments. Thus, a transition between gaseous and partially jammed states emerges without any special biotic factors other than proliferation.

**Figure 2.**
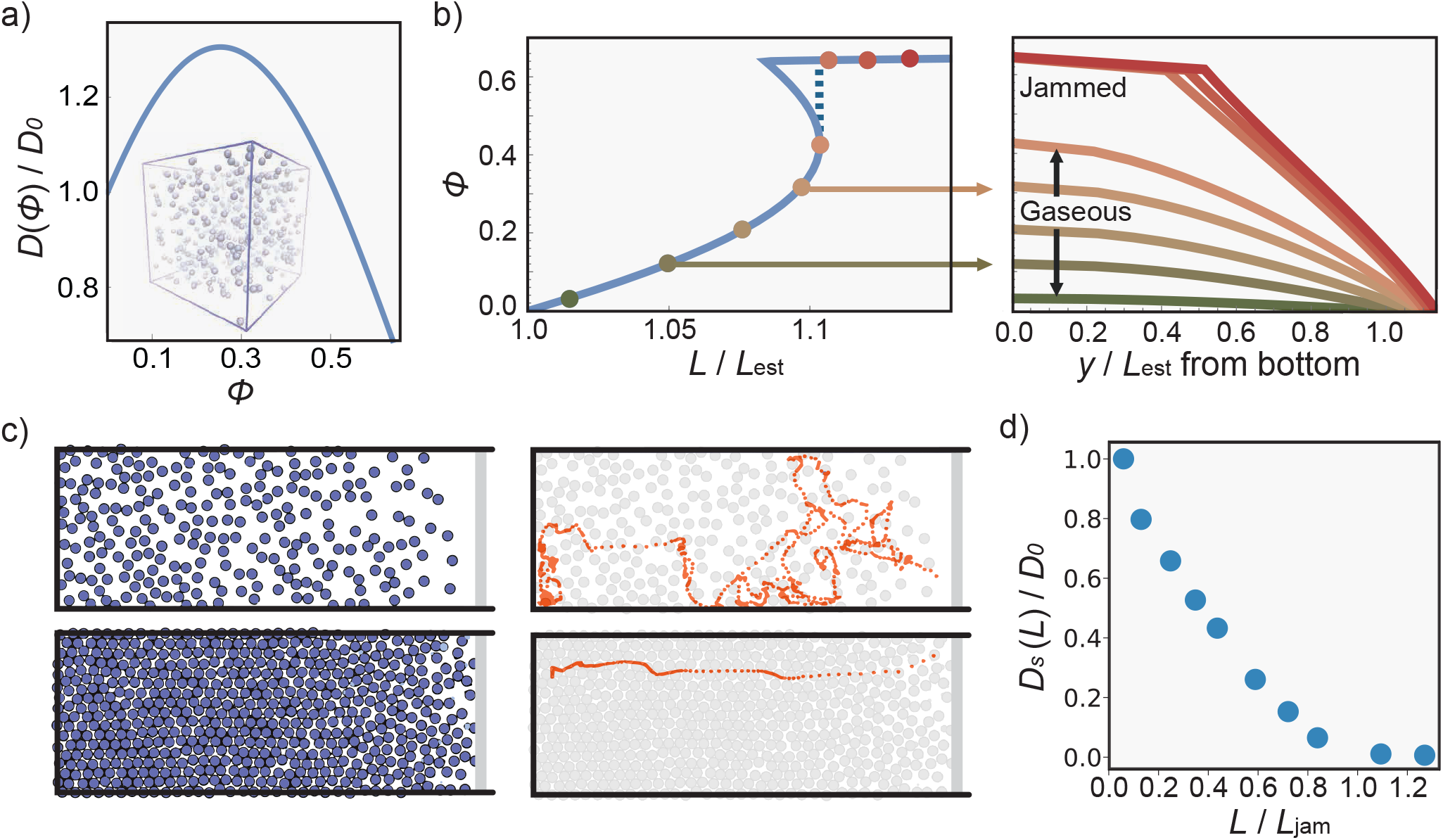
Theory: Collective motion can stabilize a growing population and drive phase separation. (a) Simulations show that the collective diffusivity of an idealized model of proliferating hard spheres in suspension is non-monotonic as a function of the packing fraction Φ = *cπσ*^3^/6 with *σ* the diameter of the particle. The negative gradients at high densities can drive a discontinuous transition towards jamming. (b) The packing fraction profile (right) was computed from the density-dependent diffusivity in (a) (see SI Sec. A.5). The maximum packing fraction (left) shows a fold bifurcation as a function of *L*/*L*_est_, resulting in a sudden transition to a (partially) jammed state. (c) Minimal simulations of proliferating soft disks and example tagged particle trajectories for gaseous *L* < *L*_jam_ (top) vs. jammed states *L* > *L*_jam_ (bottom) pores. (d) Self-diffusion, *D_s_*, in the gaseous state is larger by orders of magnitude than in the jammed state, suggesting a mechanism for an invasion barrier.

In the SI, we show that, by exploiting a mathematical analogy to a solvable Newtonian problem, the phase diagram and the density profiles (c.f. Fig. 2b for hard spheres) can be obtained exactly by numerical integration (via SI Eqs. 34 and 35) from the underlying growth and dispersal parameters. This analysis shows that the position of the tipping points depends on the entire functions *D*(*c*) and *r*(*c*) up until the tipping point and, thus, can be modulated by any means that change these functions, such as attractive interactions or quorum sensing.

Our theory also predicts that the jamming transition arises through a fold bifurcation and, therefore, should have the characteristics of a tipping point^27–29^. In particular, once a chamber becomes jammed it is not easily unjammed and requires a substantial perturbation of the control parameters (growth rate or diffusivity). This also implies that there must be a region of bistability, where in the same chamber two states are stable - one gaseous and one phase-separated state (Fig. 3a). We confirmed that, in our experiments, chambers near the jamming transition indeed show bistability (Fig. 3b and S7) by flipping from one state to another using flow modulation (Fig. S8).

**Figure 3.**
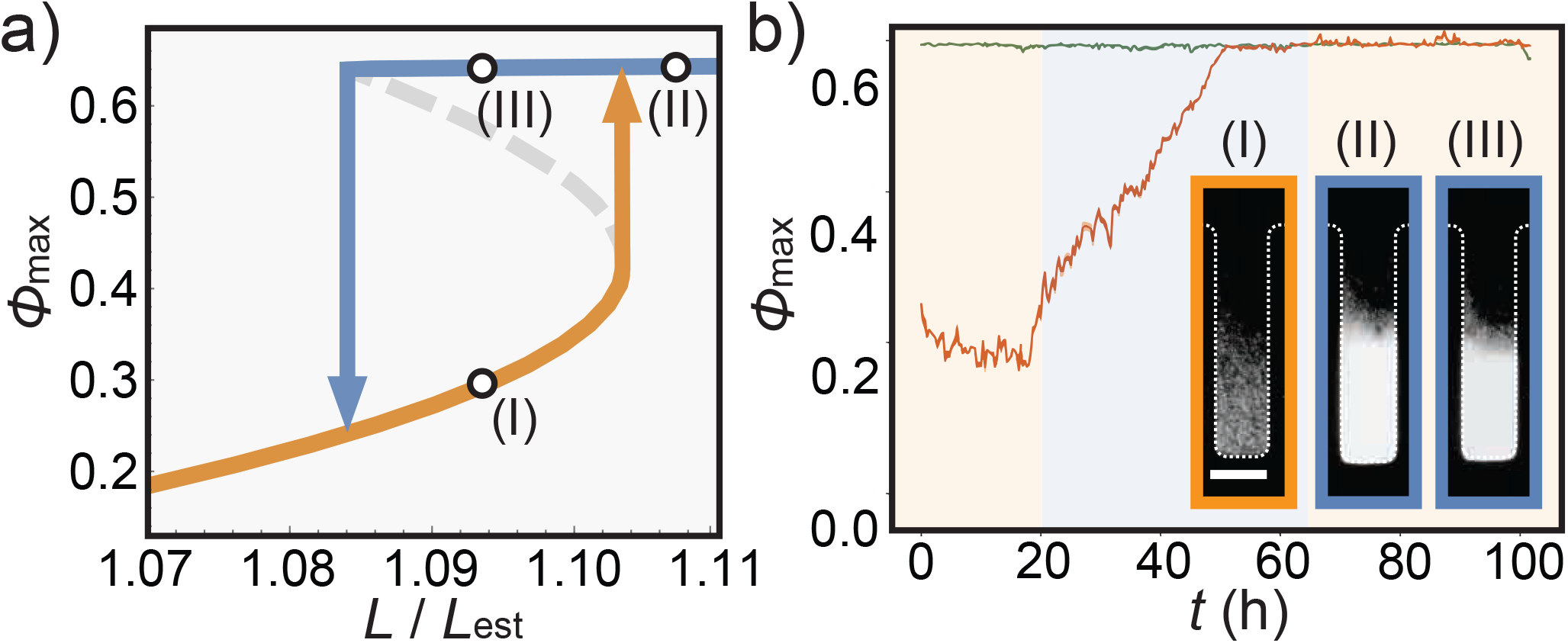
Bistability near the tipping point. (a) Phase diagram: The maximal packing fraction at steady state, Φ_max_, as predicted from the density-dependent diffusivity in Fig. 2a. When the control parameter *L*/*L*_est_ is gradually increased, the state of the system suddenly jumps from a gaseous (I) to a partially jammed state (II, arrow pointing up). If one decreases the control parameter again, the system jumps back to a gaseous state (arrow pointing down), but at a different value of the control parameter, implying a hysteresis and a region of bistability. (b) Experiments to test bistability: A flow decrease triggered in the depicted chamber the transition from gaseous (I) to jammed (II), via an effective increase of the habitat size *L*. The orange curve depicts the density increase over time. After saturation, we increased the flow again but the chamber remained in the jammed state (III) at high density (green curve). The y-axis of the plot was normalized by Φ_rcp_ ~ 0.64, random close packing of monodisperse spheres (see Fig. S7). The scale bar indicates 50 μm.

Tipping points also reveal themselves dynamically, through a dramatic slowing down near the transition point a phenomenon called critical slowing down^27^. Indeed, our time lapse Supplementary Movie S1 shows that the relaxation dynamics near the transition point becomes very slow. The smallest jammed chamber takes about 30 hours or 14 doubling times to reach steady state, compared to 6 hours or less in the largest chambers.

### Crowding-induced Drop in Diffusion

Simulations of a proliferating soft sphere model (see SI for details) further show that the cellular self-diffusion is dramatically reduced upon jamming, consistent with an onset of rigidity, except for movement of order one cell diameter per doubling induced by the division process (Fig. 2c). While in our experiments we could not track single cells in the jammed phase, we could track lineages using fluorescent tracers (Fig. S9), which also suggests self-diffusion to drop by two orders of magnitude from the gaseous to the jammed state.

A drop in self-diffusion has important consequences for species invasions. It lowers the chance for outside cells to diffusively penetrate the jammed fraction against the proliferation current coming from the floor of the chamber. Accounting for this crowding-induced diffusion barrier in a theory of strain invasion (SI Sec. C.1), we predict that the rate at which an external strain invades a jammed resident population is exponentially small in the ratio of the thickness of the jammed phase and the cell diameter. Thus, invasion of jammed populations should be an extremely rare event.

### Colonization Resistance

To test this prediction, we performed specific invasion experiments. We inoculated our device with the wild type strain of *A. indonesiensis* and waited until a steady state was reached. We then flowed in a sister strain of the same species, which was fluorescently labeled green and resistant to the drug tetracycline. Titration of tetracycline then allowed us to tune the growth rate advantage of the invading strain.

In the absence of antibiotics, we did not observe any successful invasion over the experimental time scale of five days. When we added 10 μg/mL of the antibiotic (60% of MIC), scale-dependent invasion dynamics ensued: In the initial 24 hours, the drug-sensitive populations decreased the population density (Supplementary Movie S3), thus shifting the phase boundary between gaseous and jammed to larger cavities. Over the next 48 hours, drug resistant cells entered and seized a substantial number of the gaseous chambers (Fig. 4c and Supplementary Movie 4). Upon successful invasion, the population density generally increased again. Importantly, while most of the gaseous chambers were ultimately invaded, none of the jammed chambers did (out of 7 colonized panflutes monitored over 2-5 days in 3 independent experiments). The primary effect of the antibiotic is to push the state of some of the chambers from jammed to gaseous, upon which invasion becomes possible (Fig. 4a). Thus, while crowding strongly protects jammed populations from invasion, residents can be dislodged nevertheless if they are driven past a tipping point into a more fragile (gaseous) ecological state.

**Figure 4.**
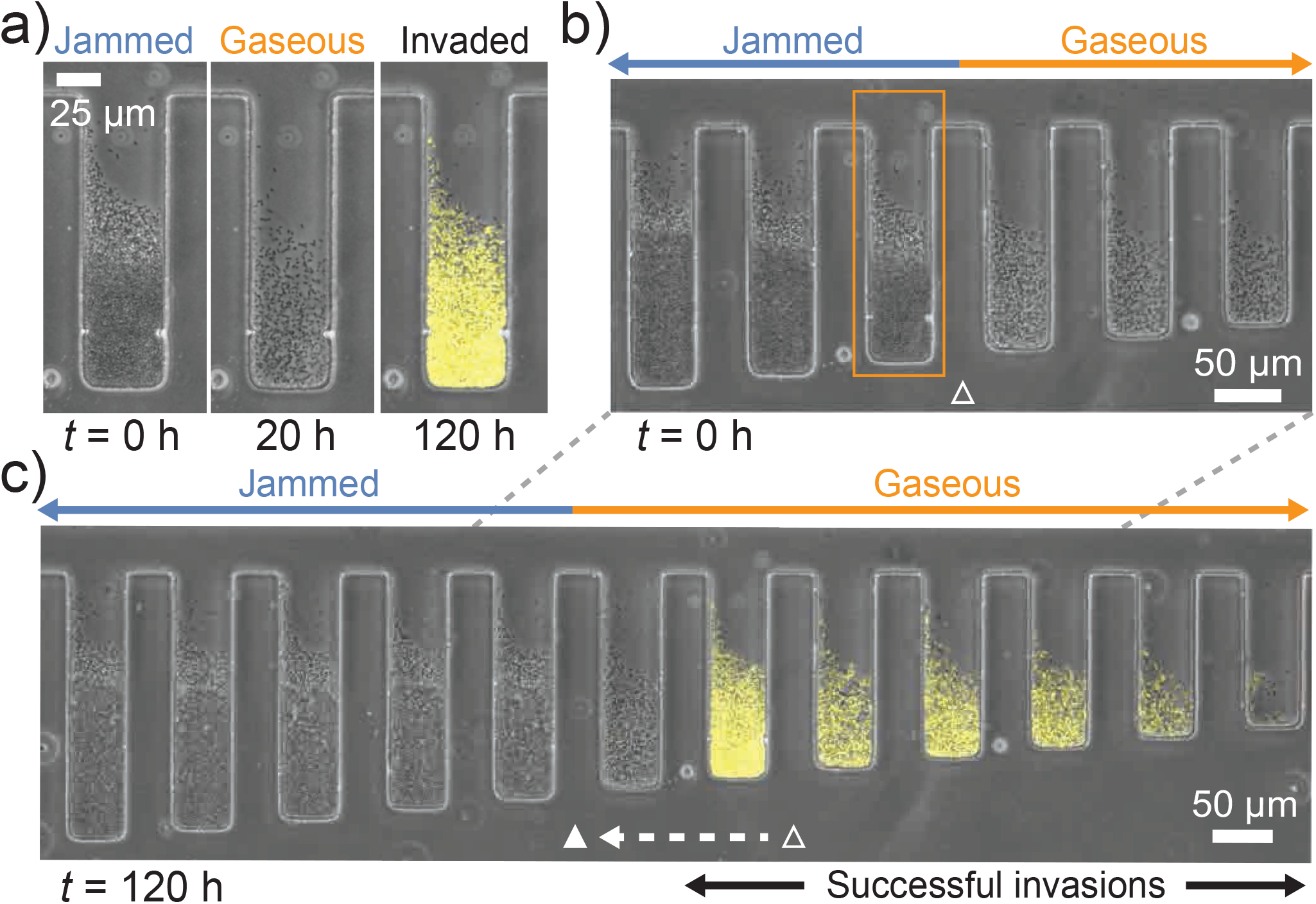
Crowding-induced colonization resistance. (a) After the chambers were pre-colonized by the wild type strain (dark), we introduced a fluorescently labeled “invader” strain (yellow). To make invasions more likely, we also increased the fitness of the invader by the simultaneous injection of an antibiotic (tetracycline) to which the invader was made resistant against. (b) A steady state of sensitive populations before the invaders were inoculated. The unfilled triangle shows the transition point between jammed and gaseous phases in the experiment. The transition was manually defined based on the bright-field darkness of the populations. (c) 120 hours after tetracycline was added to the culture medium. Drug-sensitive populations (dark) that remained jammed were not invaded. The unfilled and filled triangle show the transition points between jammed and gaseous phases at *t* = 0 h and *t* = 120 h respectively. The injection of the growth inhibitor (tetracycline) shifted the transition point.

## Discussion

We have shown that microbial colonization patterns can vary dramatically with the physical structure of their micro-environment. In particular, a crowded state with pronounced colonization resistance can arise spontaneously when the incubation scale exceeds a certain tipping point. Once pushed beyond the tipping point, it requires a substantial perturbation to break the ensuing colonization resistance, for instance by using antibiotics to trigger the reverse transition towards a gaseous phase with increased mixing (Fig. 4a).

The physical structure of the micro-environment, thus, acts as an ecological filter, permitting stable and resilient colonization by species with matching traits. By modulating the physical characteristics of this filter, hosts can actively or passively shape the pool of potential bacterial residents. Modulating endogenous micro-structures or introducing rationally-designed structures may also be considered as a strategy for precision-microbiome therapies to modulate microbial diversity.

The structure-induced stability supports the view that community assembly from potential colonizers are shaped by priority effects: whoever invades first enjoys colonization resistance against late invaders. The randomness induced by the order of strain arrival might contribute to the substantial host-to-host variability seen in some host-associated microbial communities^9, 16^.

Colonization patterns, tipping points and colonization resistance could be captured by a minimal model that accounts for growth, diffusion and leakage. This model revealed a generic fold-bifurcation generating a discontinous transition between a gaseous phase, in which cells diffuse freely, and a glassy, jammed phase. This transition differs from what is known as Motility-Induced-Phase Separation (MIPS)^26^ in the field of Active Matter^30, 31^. MIPS is associated with a spinodal instability that arises when an effective diffusivity becomes negative – an unintuitive consequence of the non-equilibrium nature of active motility^32^. In our case, the transition is triggered by the weaker condition of a (sufficiently) negative density-dependent diffusivity, which generically arises even for passively diffusing particles^33^, for example hard spheres. It would be interesting to extend our model of proliferating active matter by active motility to see how bacteria that grow and swim self-organize in confined spaces.

While the tipping points in our experiments could be explained by our minimal model, we expect that, in general, additional biotic and abiotic factors influence colonization patterns quantitatively. For example, crowding will be promoted if cells stick to one another directly or indirectly through biofilm formation, or if nutrients are supplied from the floor of the chamber. On the other hand, both establishment and jamming tend to be hindered by strong nutrient limitations or bacterial motility. While further research is needed to explore the relative importance of these factors, their impact may be anticipated theoretically using a reaction-diffusion model, which entails a flexible approach to analyze steady states (SI Sec. A.5).

More broadly, our results underscore that the lengthscale of experimentation can have a strong influence on micro-ecological processes, which could confound experiments that do not control for scale variation^34^ – a well appreciated problem in the macro-ecological context^35–37^. Flow-tunable scale variations as implemented in our Microfluidic Panflute offer a systematic experimental approach to detect or exclude scale sensitivity in culturable microbial communities. Since the time scales of microbial evolution and ecology are inter-twined, we expect such scale sensitive experiments an exciting avenue for future eco-evolutionary research^38^.

## Acknowledgments

We would like to thank all members of the Hallatschek lab for helpful discussions. We thank Carl F Schreck for useful discussion about early-stage agent-based simulations. We also thank Arolyn Conwill and Tami Lieberman for their insightful and helpful comments. The *Acetobacter* strains are generous gifts by William B Ludington, whom we also thank for useful discussions at the outset of the project. Research reported in this publication was supported by the National Institute of General Medical Sciences of the National Institutes of Health under award R01GM115851, a National Science Foundation CAREER Award (# 1555330) and the Miller Institute for Basic Research in Science. The hydrodynamics simulations were done in Molecular Graphics and Computation Facility, UC Berkeley (NIH S10OD023532). DTL was supported by NSF Grant CHE-195458.

## Data availability

All data generated and analyzed in this study are available upon request.

## Code availability

All codes used for the data analysis and the simulations in this study are available upon request.

## 1 Methods

### 1.1 Bacterial strains and culture condition

The *Acetobacter indonesiensis* strains were derived from SB003 (kindly gifted by William Ludington, Carnegie Institution for Science), which was originally isolated from lab flies (*D. melanogaster*)^10, 24^. SB003 was transformed with mGFP5 via the backbone plasmid pCM62^39^ by Benjamin Obadia^12^. For culturing, all strains were grown in MRS medium (BD) at 30 °C. Strains are selected with 15 μg/mL tetracycline (Corning Cellgro) if needed.

### 1.2 Microfluidics fabrication

The microfluidic devices were fabricated by soft lithography^40^. In order to make a master mold, a 20μm-thick layer of negative photoresist (SU8-2010, MicroChem) was spin-coated on a silicon wafer (WaferNet) and patterned by photolithography with a mask aligner (Hybralign 200, OAI) through a photomask (CAD/Art Services). On the master mold, Polydimethylsiloxane (PDMS, Sylgard 184, Dow Corning) was poured with crosslinker at 10:1 ratio and cured at 60 °C in an oven overnight. The patterned PDMS was punched to make holes for inlets and outlets. The PDMS was bonded to a glass coverslip with O_2_ plasma treatment by a reactive ion etcher (Plasma Equipment Technical Services).

### 1.3 Microfluidic cell culture

Prior to microfluidic culture, cells were streaked on a plate from frozen stock and grown in a test tube with 3 mL MRS for 1-2 days. The suspension of cells was injected into a microlfuidic device and cultured for 3-5 days with a continuous supply of the fresh medium until the system reached a steady state. The temperature was regulated at 30 °C by a microscope incubator (H201-T and UNO, Okolab), and the flow rate of the culture medium was controlled at 0.3 μL/h by syringe pumps (neMESYS, CETONI). Images were taken by inverted microscopes (IX81, Olympus. Also, Eclipse Ti, Nikon, was occasionally used for Figs. S6b and S7b.) and a confocal microscope (LSM 700, ZEISS).

### 1.4 Density profile measurement

To quantitatively measure the density profile of cellular populations in microfluidic crypts, GFP-tagged cells were cultured in a Microfluidic Panflute for about three days. After the system reached a steady state, fluorescent intensities were measured every 20 minutes for 14-48 hours. The intensities were first averaged over the horizontal direction, and then averaged over the time points at each *y*-position. They were scaled by the intensity of jammed populations to get relative cell densities. A standard error of the mean was calculated by dividing the standard deviation across time points by the square root of the approxomate number of uncorrelated time points. The latter was estimated by dividing the total duration of the time lapse movie by the typical relaxation time (6 hours) of the density profile measured in the gaseous phase (see Fig. S7d).

### 1.5 Neutral competition and invasion with fitness effect

To observe competitions of two strains with and without fitness effects, wild-type and GFP-tagged strains were co-cultured. As the GFP-tagged strain was resistant to tetracycline, with 10 μg/mL of tetracycline, the GFP-tagged cells grew normally while the wild-type strain grew slowly. We confirmed that there was no significant growth rate difference between the strains in the absence of antibiotics (Fig. S4b).

For neutral competition experiments, a 50:50 mixture of dark and GFP-tagged cells were inoculated into a Microfluidic Panflute device and cultured for two days. As each type of cells colonized chambers stochastically, we parallelized six rows of the panflutes and selected chambers with a desired initial population ratio. The population dynamics were observed with a fluorescent microscope every 20 minutes for a day.

To test the colonization resistance of jammed populations, we first cultured wild-type cells in a microfluidic device. After the populations reach a steady state, the culture medium was changed from MRS to MRS + 10 μg/mL tetracycline, and GFP-tagged cells were continuously flowed into the device. The resulting population decay and invasion dynamics were observed with a microscope every 20 minutes for two days. In addition, the snapshots of the populations were taken every day for five days.

### 1.6 Flow and temperature change experiments

To investigate the effect of the system’s parameters on the population density in microfluidic crypts, we dynamically changed the flow rate to tune the effective chamber depth. We initially cultured cells at 0.8 μL/h flow rate for three days until the system reached a steady state, and changed the flow rate to 0.3 μL/h. The decrease of the flow rate affected how deep the streamlines invaded chambers and changed the effective chamber depth by 5-10 μm (Fig. S8). After the system reached a second steady state, we recovered the flow rate to 0.8 μL/h to investigate hysteresis.

We also dynamically changed the temperature of the incubation chamber for controlling the growth rate of cells. We first cultured cells at 22 °C, where the growth rate is 0.28 h^−1^, until a steady state, and ramped up the temperature to 30 °C, where the growth rate is 0.33 h^−1^ (for the growth rate measurement, see Fig. S4d). The transition dynamics were recorded every 20 minutes with a microscope.

### 1.7 Colonization experiments with other species

Colonization dynamics in a Microfluidic Panflute were tested with various microbial species (*Escherichia coil*, *Bacillus subtilis*, *Vibrio cholerae*, *Acetobacter pasteurianus*, *Acetobacter tropicalis*, and *Lactococcus lactis*. See Table S1 for the strain details and culture media.). Cells were streaked on a plate from frozen stock, and a small number of cells from a single colony were grown in a test tube with 3 mL of a culture medium for 1-2 days at 37 °C for *Escherichia coil* and 30 °C for the other species. The cell suspension was injected into a Microlfuidic Panflute and cultured for 5-6 days with a continuous supply of fresh media until the system reached a steady state. During the microfluidic culture, the temperature was regulated at 30 °C for all species.

### 1.8 Growth rate measurement

The growth rate of cells was measured in two ways: growth assay with a plate reader and particle image velocimetry (PIV) of a jammed population on microfluidics. Prior to the measurements, cells were cultured in test tubes from single colonies for 1-2 days in MRS at 30 °C up to saturation. For the plate reader experiments, cell suspensions were diluted to 0.02 OD, and 200 μL of the suspensions were transferred to transparent flat-bottom 96-well plates (Thermo Fisher Scientific). The plates were incubated in a plate reader (Spectramax) at 30 °C, and the optical density (OD) was measured at 600 nm wavelength every 5 minutes with 30-second mixing before each measurement. The maximum growth rate was calculated by fitting an exponential curve to the initial 2-hour growth. The growth rate of *Acetobacter indonesiensis* was measured as 0.325 ±.003 h^−1^ (Fig. S4a).

For the PIV measurement on microfluidics, cells were injected into a microfluidic device and incubated in a table-top incubator until cells colonized chambers and formed stable populations. Bright-field images were taken every 3 minutes for 3 hours and analyzed with PIVlab in Matlab^41^. PIV calculated the displacements of cells per timeframe. The displacement of a cell at position *y* = *y*_0_ was caused by the growth of cells at *y* ∈ [0, *y*_0_], and therefore, the displacement at position *y* = *y*_0_ could be formulated as *d*(*y*_0_) = *y*_0_(*e*^*r*Δ*t*^ – 1). Since our timeframe (Δ*t* = 3 minutes) was much smaller than the doubling time of the cell (2.1 hours), it held *d*(*y*_0_)/Δ*t* ≈ *ry*_0_. Thus, the slope of the velocity field in the y-direction gave the growth rate. The growth rate of *Acetobacter indonesiensis* was measured as 0.332 ±.007 h^−1^ (Fig. S4d).

### 1.9 Diffusivity measurements

#### Self-diffusivity

To estimate the self-diffusivity of cells in gaseous and jammed phases, the displacement of cells was tracked over time, and the mean square displacements were calculated. A 50:50 mixture of dark and GFP-tagged cells was injected in a Microfluidic Panflute and cultured until the system reached a steady state. The motions of GFP-tagged cells were recorded with a fluorescent microscope every 30 seconds for 10 minutes for gaseous phases and every 20 minutes for 20 hours for jammed phases. The displacement of cells in gaseous phases was automatically tracked with TrackMate in Fiji^42^, and that in jammed phases was manually tracked with the Manual Tracking plugin of ImageJ.

#### Collective diffusivity

To determine the collective diffusivity, we adapted the Boltzmann-Matano analysis^43^ to the present case of a reaction-diffusion system. Under the assumption that our general reaction-diffusion model, *∂_t_c*(*y*, *t*) = *∂_y_*[*D*(*c*(*y*, *t*))*∂_y_c*(*y*, *t*)] + *rc*(*y*, *t*), is valid, we can express the density-dependent diffusivity *D*(*c*) in terms of the steady state density profile as follows:

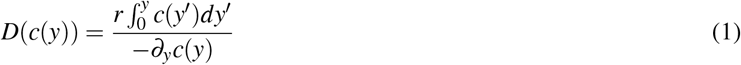

This equation allows us to estimate *D*(*c*) from the exponential growth rate and the steady state density profiles. The steady-state density profiles were determined from the temporal average of the fluorescent intensity of time lapse movies. For the data in Fig. S2c, we averaged the density profile over 20 frames (7 hours) and locally approximated it with a quadratic function by the Savitzky–Golay method^44^ to extract *∂_y_c*(*y*). We excluded the y-region 20% from the opening where the flow impacted the tail of the density profile, and excluded the y-region 20% from the bottom where (*∂_y_c*(*y*))^−1^ was diverging.

### 1.10 Fluid dynamics simulations

The fluid dynamics of the culture medium flow through the cavity structures of our microfluidic devices were simulated using COMSOL. As a simple geometry, we defined a 500 μm × 50 μm × 20 μm supply channel with a 50 μm × 150 μm × 20 μm cavity in the middle. The fluid dynamics was modeled as incompressible Stokes flow subject to no-slip boundary conditions at the walls and a constant flow rate. To see how the flow field depends on external control parameters, we varied the depth of the cavity (30-150 μm) and the flow rate (100 μm/s and 250 μm/s mean flow rate).

## 2 Descriptions of Supplementary movies

- S1: Jamming dynamics and gaseous phases. The movie shows the lengthscale-dependent colonization in a Microfluidic Panflute over 35 hours. While cell populations get jammed in long chambers, they remain gaseous in short chambers. The approach to the steady state takes markedly longer in chambers close to the transition, which is characteristic of critical slowing down near tipping points.
- S2: Formation of a jammed shockwave. The movie shows the movement of the jamming front over 20 hours. The culture medium flow came from the top to the bottom.
- S3: Population decay upon drug injection. The movie shows the decay of a drug-sensitive population over 14 hours upon injection of 10 μg/mL tetracycline. The sharp decrease of the growth rate of cells results in the phase transition from initially jammed to gaseous.
- S4: Invasion of an advantageous strain. The movie shows how a drug-sensitive resident population (dark) is invaded by drug-resistant cells (yellow). The fitness of the established drug-sensitive bacteria are suppressed by the addition of 10 μg/mL tetracycline. Successful invasions are observed 21 hours after the injection of the drug and the advantageous strain.

## Supplementary Information

### A Theory of colonization

Here, we describe our modeling approach that helps us to interpret and predict the relaxation and steady-state properties of the bacterial populations in our variable length cavities. We begin by describing a general mathematical framework applicable to a wide range of situations. We then restrict the model to our specific experimental setup, which allows us to make a number of simplifying assumptions.

#### A.1 General reaction-diffusion model

In order to describe the combination of growth and cell movement, we employ a reaction-diffusion model for the packing fraction Φ(**r**, *t*) at position **r** and time *t*, which is the concentration multiplied by the cell volume. In general, the rate of change of the packing fraction is given by

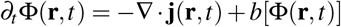

in terms of the divergence **∇**·**j** of a flux **j**, a three dimensional vector, and the cell production rate *b*[Φ], which depends on the packing fraction itself, for exampled *b*[Φ] = *rφ* in the case of exponential growth with rate *r*.

The flux **j** describes the magnitude and direction of the flow of cells and can be expressed as

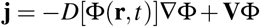

in terms of the cell diffusivity *D*, the gradient **∇**Φ of the packing fraction and an advection velocity **V**, a three dimensional vector.

#### A.2 Context-specific simplifications

A number of simplifications can be made due to the specific setup of our experiments. Here, we describe these simplifications, and note under which conditions they might break.

- *Diffusion is purely passive*. The bacteria in our experiments have no motility mechanism and, therefore, diffuse just like passive particles under the influence of thermal collisions with the solvent particles. However, many bacteria have flagella allowing them to swim and perform chemotaxis to chase the source of certain chemical cues. Although the behavior of swimming bacteria can be quite complex, they are often well described by an advection term to describe chemotaxis and an effective diffusivity, which has a non-trivial motility-induced density-dependence^26^.
- *Advection is absent*. In other systems, advection may have to be incorporated to describe chemotaxis (pervious point) or the influence of gravity or fluid flow. Fluid flow is a particularly important aspect in the lumen of the gut^45^ but also can also arise in microscopic pores, for instance, in skin pores when sebum is exuded^16^.
- *The setup is effectively one-dimensional*. The concentration profile in the narrow crypts of our panflute device is approximately uniform along the directions perpendicular to the symmetry axis of the crypts. This allows us to restrict our discussion to the dynamics along the symmetry axis – the *y*–direction. For wide or high crypts, a three-dimensional description is necessary, especially when hydrodynamic instabilities drive motion in the direction perpendicular to *y*. We expect such higher-dimensional dynamics to be a fruitful topic for future work, in particular because it cannot be mapped to an effective Newtonian dynamics (described below).

Under these simplifying assumptions, which match our experimental conditions, we obtain an effectively one-dimensional reaction-diffusion model that reads

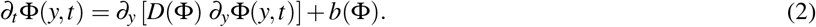

The boundary conditions are fixed by demanding that cells shall not exit through the floor of the chamber, *∂_y_*Φ = 0, and that the density vanishes at position *L*, Φ(*L*, *t*) = 0.

#### A.3 Linear Stability Analysis

In order to determine the onset of population growth in our microfluidic crypts, we have to determine the conditions for which an empty crypt is stable against an inoculation with cells. To this end, we perform a linear stability analysis of our model in the low density limit.

The linearized reaction-diffusion equation Eq. 2 reads

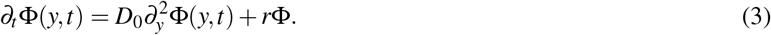

in terms of the low-density diffusivity *D*_0_ and the constant growth rate *r*.

We first expand the cell density as

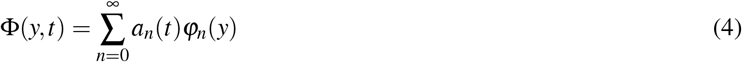

in terms of cosine modes

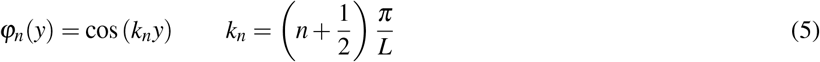

which are orthonormal

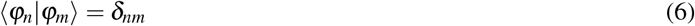

with respect to the scalar product

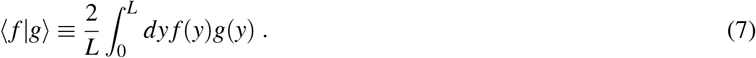

Inserting the expansion Eq. 4 into the linearized reaction-diffusion equation Eq. 3 and then projecting onto the normal modes yields simple amplitude equations,

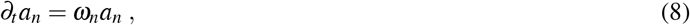

where the frequency *ω_n_* of the *n*^th^ mode is given by

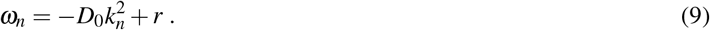

The mode amplitudes *a_n_* therefore obey *a_n_* = *a_n_*(0) exp(*ω_n_t*) with the prefactors fixed by the initial conditions.

For the empty state to be linearly stable, we require all *ω_n_* to be negative, meaning that all mode amplitudes exponentially decay to zero. As the slowest growing mode is *n* = 0, this implies 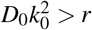, or

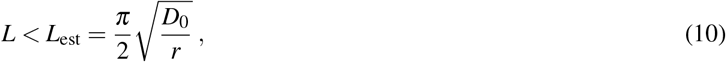

revealing the establishment transition discussed in the main text.

Note that the density decay of non-growing particles (*r* = 0), which merely diffuse out of the chamber, is on long times controlled by the slowest decaying mode, 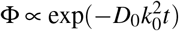. Since, at the establishment transition, we have 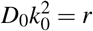, we see that the diffusive “half-life” of the bacteria just equals their doubling time. This confirms the intuition that the establishment transition occurs when the diffusive outflow is balanced by growth.

#### A.4 Steady state at low packing fraction

To explore the nature of the gaseous phase, it is useful to study the limit of low packing fractions, where the diffusivity takes the form^33^

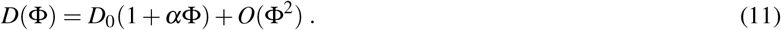

The numerical coefficient *α* measures the leading order change in diffusivity with increasing packing fraction and depends on the shape of the particles as well as their interactions. For repulsive particles, such as our main model system *Acetobacter indonesiensis*, *α* is positive. Detailed analytical results are available for hard spheres, yielding *α* = 1.45^46^. Strong attraction can lead to negative *α*^33^.

When we include the non-linear packing fraction dependence to leading order, we obtain the following equation of motion

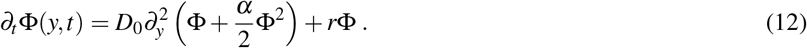

If we expand Φ(*y*, *t*) in terms of the normal modes as in Eq. 5, we find

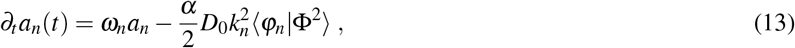

where 〈*φ_n_*|Φ^2^) is the projection of Φ^2^(*y*, *t*) on to the *n*^th^ mode.

We expect the small density approximation, Eq. 13, to be appropriate when *L* just slightly exceeds *L*_est_. Thus, we can introduce the small parameter

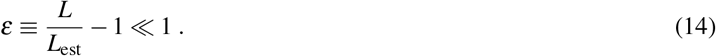

Our discussion of the establishment transition has shown that the frequency of the lowest mode vanishes right at the transition, *ω*_0_ = 0, and that the frequencies of all other modes is finite, i.e. *ω_n_* = *O*(1).

For *ε* small but finite, we still have that the higher modes have linear relaxation frequencies of order one, *ω_n_* = *O*(1) for *n* > 1, but the frequency of the lowest mode now assumes small positive frequency of order *ε*,

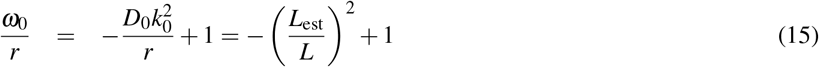

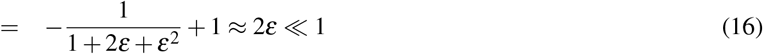

Combining *ω*_0_ =O(*ε*), *ω*_*n*>1_=O(1) with Eq. 13 shows that, at steady state, *a*_*n*>1_ with *n* > 1 is of higher order in *ε* than *a*_0_.

Thus, to leading order in *ε*, we have Φ(*y*, *t*) = *a*_0_*φ*_0_(*y*, *t*) + *O*(*ε*^2^), which simplifies the amplitude equation Eq. 13 to

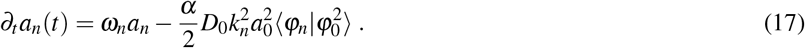

The scalar product on the right hand side evaluates in general to

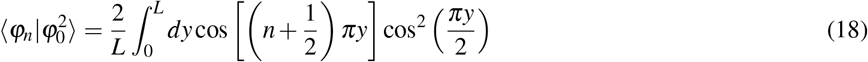

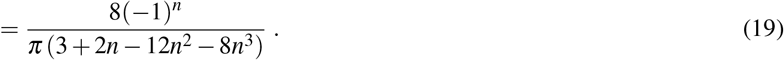

As before, we confine our attention to the slowest growing mode, *n* = 0, for which 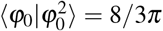 leading to the closed amplitude equation,

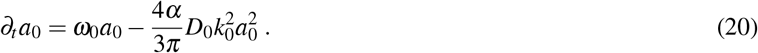

the steady state density will be small allowing us to expand

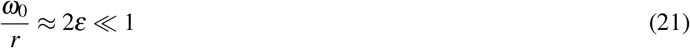

in terms of

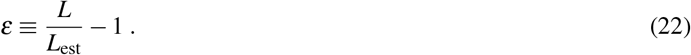

Rescaling time *τ* = *rt* and using the leading order approximations 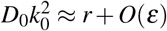 and *ω*/*r* ≈ 2*ε* (Eq. 16), we obtain

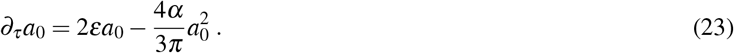

For the inoculation of an initially empty chamber, this logistic differential equation yields the simple prediction

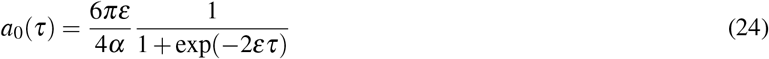

up to a shift in time.

To illustrate these results, we consider the example of hard spheres for which *α* = 1.45 is known exactly (from Eq. (6.12) in Ref.^46^). Our lowest order expansion Eq. 24 then predicts a packing fraction at the floor of the chamber of *c*(*y* = 0) = *a*_0_ cos(0) = *a*_0_ = *ε*6*π*/(4*α*) ≈ 3.25*ε* at steady state (*t* → ∞). Extrapolating from this lowest order expansion, we may estimate that a chamber length of no more than *ε_j_* = 24% above the establishment length is needed for jamming to occur, because then the maximal density at the floor, 3.25*ε_j_* = 0.64, approaches random close packing (Φ_rcp_ ≈64% for monodisperse spheres). This simple estimate of course ignores non-linear feedbacks, which typically leads to an earlier onset of jamming, as seen in Figs. 2b and 3a.

Of particular significance is also the predicted relaxation time to the steady state, the inverse of which is often taken as a measure for the resilience of an ecological system^47^. From the exponent in Eq. 24, we see that this relaxation time is given by 1 /(2*εr*), which is independent of *α* and, notably, diverges near the establishment transition. Thus, relaxation can take long — much longer than the diffusive exploration of the chamber, which takes about one cell doubling near the establishment transition (the diffusive half-life of particles in a crypt at establishment length just equals the doubling time, see Sec.A.3). Therefore, we generically expect a time scale separation between relaxation of the density profile towards the cosine shape (fast) and relaxation of the amplitude of the cosine (slow).

Although the cells in our experiments are neither spherical nor monodisperse, we expect the above results to apply up to pre-factors. For example, since *ε* remains small up to the jamming transition also in our experiments, one would expect long relaxation times throughout the gaseous phase. Indeed, relaxation in our flow shift experiments took up to 10 h or five doublings (see green curve in Fig. S7).

#### A.5 Mechanical analogy predicts steady-state colonization patterns

Let us consider the one-dimensional reaction-diffusion equation for the cell packing fraction Φ(*y*, *t*) at position *y* and time *t*

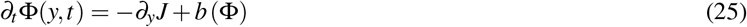

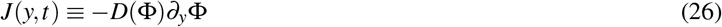

where we allow for an arbitrary density-dependence in both the collective diffusivity *D*(Φ) as well as the growth rate *b*(Φ). As pointed out in the main text, the linear growth rate in our experimental system is to a good approximation constant, which corresponds to *b*(Φ) = *r*Φ. The mathematical treatment in this section is independent from this simplifying condition.

We define the quantity

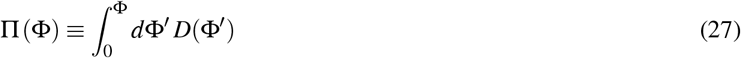

such that the diffusive current is given by

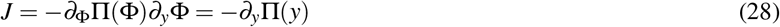

where we identified П(*y*) ≡ П[Φ(*y*)] to simplify the notation. Eq. 28 formally implies that a gradient of Π(*y*) drives a current just like a conventional pressure gradient would. We therefore call **Π**effective pressure. Since for passive diffusion^2^, we must have *∂*_Φ_П(Φ) = *D*(Φ) > 0, we can invert the equation of state to obtain a the packing fraction Φ(П) as function of effective pressure П.

Next, combining Eqs. 25, 27 and 28 yields at steady state

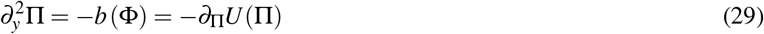

where we defined an effective potential

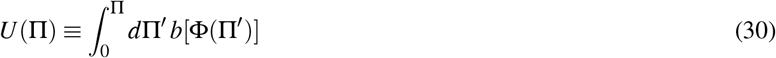

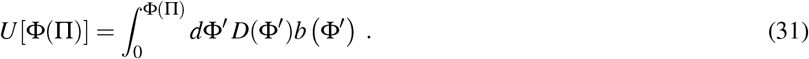

The boundary conditions imply П(*L*) =0, manifestly ensured through Eq.27, and *∂_y_*П(*y* = 0) = 0, which we will account for below.

Notice that Eq. 29 is formally identical to Newton’s equation for a particle at position П(*y*) at time y freely falling in a potential *U* (П). Hence, we can use the principle of mechanical energy conservation to immediately predict the velocity *∂_y_*П of the moving particle in our mechanical analogy, which in our reaction-diffusion problem corresponds to the negative particle current

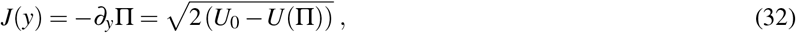

manifestly satisfying the no-flux boundary condition *∂_y_*П(*y* = 0) =0. Note that we use the notation *U*_0_ ≡ *U*(П_0_) and П_0_ ≡ П(*y* = 0).

Integrating Eq. 32 over the *y* yields

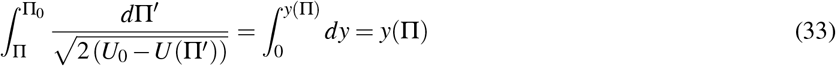

or

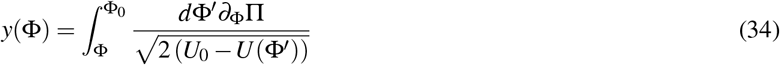

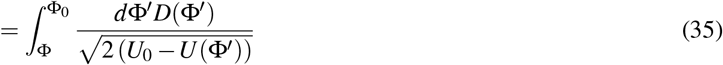

where *U*(Φ) = *U*[Φ(П)], as defined in Eq. 31. Calculating *y*(0) = *L* gives us the effective chamber length given a maximal packing fraction Φ_0_ at the floor (*y* =0) of the chamber, which is how we determined Figs. 2b and 3a. Since the function *y*(Φ) is monotonous, it can also be inverted to determine the position-dependent packing fraction Φ(*y*), shown in Fig. 2b for hard sphere,again given the maximal packing fraction Φ_0_ at the floor of the chamber.

The integrals in Eqs. 35 and 31 can be solved numerically without problems for any *D*(Φ) > 0 and *b*(Φ), provided *U*(Φ) < *U*_0_ along the trajectory.

#### A.6 General approach to compute the phase diagram

Generically, the shape of the potential *U*(Φ) will start at *U*(Φ=0) =0 and increase monotonically because *U*′ = −*D*(Φ)*b*(Φ) ≤ 0, unless we allow for a region of negative net growth *b* < 0. The behavior near Φ = 0 is quadratic since *U*′(Φ = 0) = 0 from the no-flux boundary condition and we assume analyticity of *D* and *b*. For small Φ, we expect *D*′ > 0 suggesting that the potential at first rises faster than a parabola. At larger Φ we expect, instead, a negative slope of *D*(Φ) resulting in a flattening of the potential until jamming leads to rapid rise of the potential.

To determine the density at the floor of the chamber, we have to solve the following problem: Let a point mass move down this energy landscape from Φ = Φ_0_ back down to Φ = 0. The time it takes the moving mass to reach Φ = 0 has to equal the length *L* of the chamber. If the sojourn time is too small (large) we have to increase (decrease) Φ_0_. Mathematically, we can formulate this condition using Eq. 35,

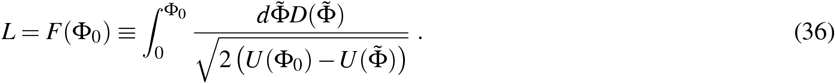

We have thus obtained an equation that can be used to determine a phase diagram as in Fig. 3a from any *D*(Φ) and *b*(Φ). Multiple steady states exist if there are multiple Φ_0_ values with *identical* sojourn times. The trivial case for which this happens is a simple parabola, which corresponds to the case without density-dependence, *D* = const. and *b*(Φ) = *r*Φ. Then, we have a critical length where any density Φ_0_ will lead to a marginally stable steady state.

More realistically, multiple steady states occur if the potential fluctuates around a parabola. The density interval supporting multiple steady states is bounded by Φ_0_-values for which

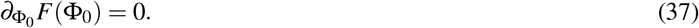

This condition can be computed explicitly as follows,

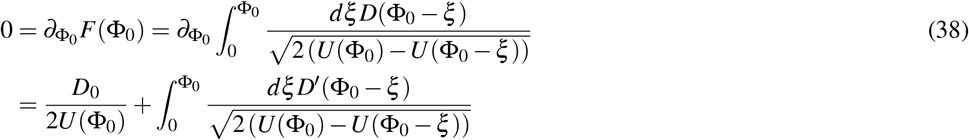

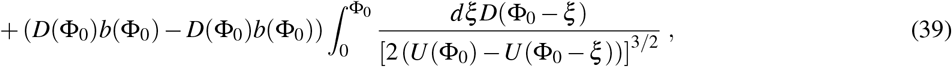

where we substituted 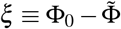 in the first line and using Eq. 31 to express the derivative of *U* in the last line. If Eq. 39 has any solution there must exist multiple steady states.

#### A.7 Collective diffusion for hard spheres

In general, *D*(Φ) is a collective diffusion coefficient as opposed to a tracer diffusivity or self-diffusion coefficient. With purely passive diffusion, *D*(Φ) can be decomposed into the product *D*(Φ) = *μ*(Φ)*k_B_T∂_Φ_P*(Φ) of two terms with an intuitive interpretation:

- *μ*(Φ) is the collective mobility, a transport coefficient that describes the sedimentation velocity and is typically a decreasing function of packing fraction Φ. As a transport coefficient it depends on the equations of motion, and the treatment of hydrodynamics.
- *P*(Φ) is the osmotic pressure of the cell suspension, and its derivative is proportional to the (osmotic) bulk modulus, which has to be positive. Because *P*(Φ) is a pure equilibrium quantity, it can be readily obtained from Monte Carlo simulations without modeling the surrounding fluid at all.

For a system of hard spheres, a multipole expansion of the effective hydrodynamic interaction^48^ can be used to extract the collective mobility for particles immersed in an incompressible fluid. With collective mobility computed using this method, we confirmed the empirical Richardson-Zaki scaling form^49^, *μ*(Φ) = *μ*_0_(1 – Φ)^η^ where *μ*_0_ is the particle mobility in the dilute limit. We extracted *η* = 5.8 from a linear regression of *μ*(Φ). The fit and accompanying data are shown in Fig. S11, which over the range of packing fractions considered are in good agreement. Similarly, the equation of state of hard spheres is known to be well described by a Carnahan-Starling equation^50^. In terms of the packing fraction, the pressure *P*(Φ) is given by, *P*(Φ)/*k_B_T* = (6Φ/*π*)(1 + Φ + Φ^2^ – Φ^3^)/(1 – Φ)^3^. Taken together, we obtain the collective diffusion coefficient via *D*(Φ) = *k_B_Tμ*(Φ)*∂*_Φ_*P*(Φ), which was used together with *b*(Φ) = *r*Φ to determine the hard sphere phase diagram Fig. 3a in the main text using Eq. 36.

It is worth noting that that the resulting collective diffusion coefficient for hard spheres is merely an approximation designed to capture the non-linear behavior for modest to high packing fractions. Exact results are often available for the linear expansion coefficient *α* of the diffusivity *D*/*D*_0_ ≈ 1 + *α*Φ+h.o.t. (see e.g. Ref.^33^), valid at low density, which can be useful for estimating the behavior of the gaseous phase at low densities. For instance, in Sec. A.4, we used the exact result *α* =1.45^46^ for hard spheres to estimate the steady state density, rather than *α*_approx_ = 8 – *η* from the above approximation Richardson-Zaki/ Carnahan-Starling approximation.

### B Proliferating soft disk simulations

To see which phase transitions emerge in a minimal model that only includes proliferation, cell diffusion and cell repulsion, we explicitly simulate a mechanical system of soft disks that undergo Brownian motion and that divide with a constant rate.

#### B.1 Model details

Each particle, *i*, obeys a stochastic equation of motion for its position in two dimensions, ***r**_i_* = {*x_i_*, *y_i_*},

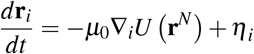

where *μ*_0_ is the time single particle mobility resulting from the surrounding fluid, *η_i_* is a Gaussian random variable with mean 〈*η_i_*〉 = 0 and variance 〈*η_i_*(*t*)⊗*η_j_*(*t*′)〉 = 2*k_B_Tμ*_0_1*δ_ij_δ*(*t* – *t*′) where *k_B_T* is Boltzmann’s constant times the temperature of the fluid and is diagonal for each particle and cartesian component. We solve this equation using a standard first order Euler discretization. In addition to Brownian motion, the particles move in response to a potential *U*(***r**^N^*) that depends on the full configuration of the system, denoted **r**^*N*^. At any time, there are *N* total particles, with *N*_m_ maturated mother particles and *N*_d_ growing daughter particles. The interaction potential is decomposable into *U*(**r**^*N*^) = *U*_b_ (**r**^*N*^) + *U*_r_ (**r**^*N*^) + *U*_w_ (**r**^*N*^) where *U*_b_ is the bonding potential between mother and daughter particles, *U*_r_ is a purely repulsive interparticle interaction and *U*_w_ is a confining potential.

The bonding potential is taken to taken to be a simple Hookian spring,

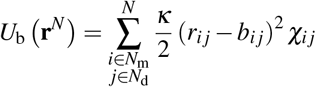

with stiffness *κ*, rest length *b_ij_*, and *χ_ij_* = 1 if particles *i* and *j* are a mother daughter pair and *χ_ij_* =0 otherwise. The specific form of the repulsive interparticle potential is taken to be pairwise decomposable

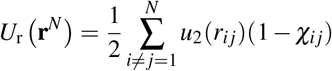

where the pair potential *u*_2_(*r_ij_*), is a WCA potential^51^

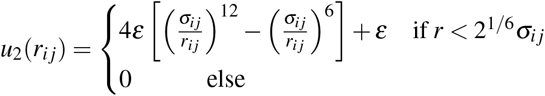

where *ε* is a characteristic energy scale and

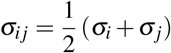

where *σ_i_* is the characteristic size of particle *i*. All pairs of particles, excluding bonded mother and daughter pairs, interact with these are excluded volume interactions. Finally, the confining potential restricts the the particles motion to approximately an area *A* = *L_x_L_y_* by imposing a steep potential at *x* = 0, *x* = *L_x_*, and *y* = 0. Specifically, the external wall potential has the form,

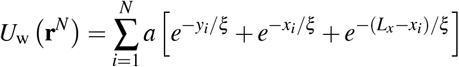

with *a* the amplitude of the confining potential and *ξ* its characteristic length scale.

The mechanical system outlined above is conservative and describes the motion of a collection of overdamped particles with a simplified, local description of hydrodynamics^52^. At long times, absent added external forces, its evolution would be consistent with thermal distribution and would conserve particle number. In order to model the growth of the bacterial population and division of an individual mother daughter pair, we endow the daughter particles with a time dependent effective size through the deterministic equation of motion

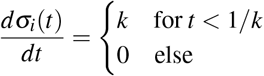

with boundary condition *σ_i_*(0) = 0 and growth rate *k*. Similarly, to model the budding of the daughter from the mother, the rest distance *b_ij_* changes in time with the deterministic equation of motion

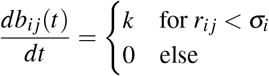

with boundary condition *b_ij_*(0) = 0 and the same growth rate *k*. When then impose that when the distance between the mother-daughter pair *r_ij_*(*t*) has grown past the effective size of the mother particle, *σ_i_*, we sever the bond potential by *χ_ij_* → 0 between mother-daughter pair *ij*, provided *t* > 1/*k*. An illustration of this criteria is shown in Fig. S12. This growth rule results in an increasing excluded area of the mother-daughter pair that grows approximately linearly in time. All of the results in the main text and below employ this rule with a fixed *k*. Growth rules that affect in an exponential increase in the excluded area with time yield qualitatively similar results. Further the results presented are for a fixed growth rate *k*, but generalizations for populations evolving with a distribution of growth rates are also qualitatively similar, provided the distribution is narrow. Synchronized with the bond breaking event, we add new daughter sites to each of the newly divided particles and reinitialize the size and rest length to 0 for each of them. This last step breaks particle number conservation. An illustration of the subsequent increase in *N*_m_ is also shown in Fig. S12, which absent mitigating factors will grow exponentially in time.

Finally, consistent with the experiments, we apply an absorbing boundary condition at *y* = *L_y_*. The absorbing boundary condition and particle number growth balance at steady-state, resulting in a mean particle number that depends on the geometry and model parameters. We adopt a unit system where *μ* = *ε* = *k_B_T* = 1, and mature particles sizes *σ* = 1. This implies that lengths are defined in multiples of the mature particles size, *x* → *x*/*σ* and times in units of the diffusion time for an isolated particle to move its diameter, with *D*_0_ = *k_B_Tμ*_0_ and accompanying units of time, *t* → *t*/(*D*_0_*σ*^2^). The stiffness *κ* in principle allows for mechanocoupling between the division and local packing environment, with deterministic division occurring when *κ* is much larger then the local stress on the mother-daughter pair. In this work, we consider this limit and take *κ* = 100. The confining potential parameters are taken as *a* = 10 and *ξ* = 1.

#### B.2 Nonequilibrium phase diagram

We have studied the particle based model described above and have found qualitative agreement with both the experimental results and simplified theory presented in the main text. Specifically, we have studied a system with confinement defined by fixed spatial scale *L_x_* = 15 and *L_y_* = 40, seeded with an *N*_m_ = 10 initial mother particles. We have found that 10 division cycles is sufficient to relax to a steady state density in the chamber, and evaluated expectation values by averaging over a minimum of 30 additional division cycles. Further, 3-5 independent simulations are used for each expectation value reported. Studies of the effective reaction diffusion model suggests that the function dependence of the system on the length of the chamber enters relative to the establishment length 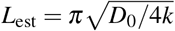, so rather then studying different *L_y_*’s, we fixed *L_y_* = *L* and study the dependence on the division rate *k*, and thus *L*_est_.

First, we studied the stead-state density distribution in the chamber. For *L*/*L*_est_ < 1, as expected the density in the chamber is 0 at steady-state on average. For *L*/*L*_est_ > 1, the particles are able to colonize the chamber, and evolve a stationary density distribution. The distribution can be characterized analogously through local packing fraction Φ(*y*) = *ρ* (*y*)*π*/4 where 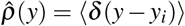. Generally, the discrete size of the particles relative to a flat wall will result in an oscillatory density profile when the overall density is larger, *ρ* > 0.1, which is a result of density correlations induced by their excluded area. Such an oscillatory density profile is not predicted by the simple reaction diffusion model employed in the main text. In order to make contact with that perspective, we report in Fig. S13 density profiles coarse-grained over a the length-scale of the particle^53^. We achieve this using by convoluting the number density with a Gaussian,

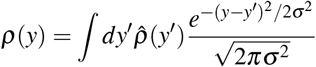

which smooths the profile out. Further, we consider the contribution of the density only from the mother particles, and evaluate expectation values for times that are integer multiples of 1/*k*.

Fig. S13 specifically reports conditions for *L*/*L*_est_ = {1.2,2,4}. We also compare those calculations to the predictions of the reaction diffusion model. For *L_y_*/*L_est_* = 1.2, the density is small enough that we find good agreement with the predicted cosine profile, Φ(*y*) = Φ(0)cos(*πy*/2*L_y_*). At elevated *L*/*L*_est_, deviations of the cosine profile are found and specifically at *L_y_*/*L*_est_ = 4, the distribution is flat in the interior of the chamber with an exponential boundary layer that brings the packing fraction to 0 at *y* = *L*. Using a parameterization of the collective diffusion constant *D*(Φ), evaluated by computing the packing fraction dependent mobility *μ*(*φ*) that is well described by *μ*(*φ*) ≈ *μ*_0_exp(−1.70Φ – 0.18Φ/(1 - 1.33Φ)), and the equation of state well described by *P*(Φ)/*k_B_T* ≈ 1.27Φ + 2.55Φ^2^ – 9.35Φ^3^ +42.72Φ^4^ of the WCA disks, consistent with previous estimations,^54, 55^ we can numerically solve the reaction diffusion equation and determine the packing fraction profile for *L*/*L*_est_ =4. This predicted profile is in good agreement with the coarse-grained profile from the simulations.

To estimate the boundaries between the extinct, established, and jammed phases, we have computed the coarse-grained value of the packing fraction at the wall as a function of *L*/*L*_est_. This is shown in Fig. S14. As anticipated by the reaction diffusion analysis, the establishment transition occurs for *L*/*L*_est_ ≈ 1, which is consistent with our findings that *L* < *L*_est_ the chamber is empty at steady-state. For *L* > *L*_est_ the maximum packing fraction gradually increases until *L* = *L*_jam_ ≈ 3.3 where for this two-dimensional system we find an abrupt change in the maximum packing fraction. The amplitude of this change is small, reflecting the small change in the density upon freezing of the WCA disks, which is around 2%^56^. Indeed, the pressure measured at the wall at *y* = 0, computed from the average force per unit length of wall, 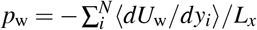, surpasses the coexistence pressure for the freezing of WCA spheres at this value of *L*. As illustrated in Fig. 2 in the main text, for *L* > *L*_jam_ the system exhibits noticeable crystallinity.

#### B.3 Self diffusion calculations

In order to estimate the self-diffusion coefficient in the chamber as a function of *L*/*L*_est_, we consider only the diffusivity in the *x* direction, the direction orthogonal to the open end. This is because there exists a net mass current in the *y* direction, so motion is convective in that direction rather than diffusive. For diffusion in the *x*, we use the standard definition

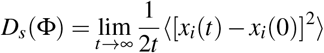

as the long time limit of the mean squared displacement. However two difficulties arise in applying this relation to extract the self diffusion. First, particles leave the chamber at a finite rate, due to the absorbing boundary condition at *y* = *L*. This requires that we average particles’ displacements only up to their lifetime in the chamber. In order to gather sufficient statistics and increase the lifetime of individual particles, we use a larger chamber *L_y_* = *L* = 80. Second, the finite size of the chamber in the *x* direction, *L_x_* means that particles will only exhibit diffusive dynamics for mean squared displacements 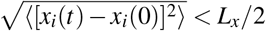, and thus the long time limit strictly goes to 0. We find using *L_x_* = 30 to be large enough that there exists a sufficient separation of timescales between the onset of diffusive motion and its reduction due to confinement that we easily extract a pseudo-time independent self-diffusion constant. This data is reported in Fig. 2.

### C Theory of Invasion

For a foreign strain to invade a pre-occupied chamber, an invader has to overcome two hurdles: #1) *Infiltration*: the invader diffuses from the outside against the gradient to reach a favorable position near the floor of the chamber. #2) *Take over:*the invader’s descendants displace the resident population through a combination of chance (genetic drift) and competitive advantage.

Our experimental observations in combination with our simulations indicate that the main reason for the colonization resistance is the difficulty for outside cells to diffusively penetrate chambers that are already filled. Infiltration is rare even in gaseous chambers, but nearly impossible in jammed populations where a narrow strip of founder cells at the floor of the chamber is insulated from the outside by diffusion barrier of descendant cells. Accordingly, the biggest impact of the antibiotic occurs in cavities that, due to the growth rate detriment of the resident, become unjammed and hence invadable (Fig. 4). The larger the growth rate detriment, the larger the range of chamber length that are driven across the tipping point.

In the limit of weak selection, and large chamber population size, infiltration happens on a much faster time scale than take over. This allows to analyze the dynamics in two separate steps. The second, takeover, step is familiar from well-mixed populations. The rate of successful take over depends on the competition between selective advantage of the faster growing invader and genetic drift. The first challenge, however, uniquely depends on the spatial structure of the colonized cavities, which is why we mainly focus on the infiltration step.

The infiltration step is best analyzed backward in time. As we follow the lineage of a randomly chosen cell backward in time, it is advected towards the floor of the chamber, reflecting the intuitive location advantage discussed above. A balance between self-diffusion and advection leads to a steady state lineage distribution, whose extent scales as the ratio of self-diffusivity and advection velocity. While the advection velocity is very similar between gaseous and jammed populations, self-diffusion differs dramatically, by four orders of magnitude, compressing the ancestor distribution in the jammed population to just few cell layers at floor of the chamber.

Our mathematical analysis in the next section shows that the neutral invasion success of an injected invader is proportional to 1/*N* times the ratio of the invader density in the supply channel and the mean density of the population. This shows that gaseous population do enjoy some colonization resistance, compared to well-mixed populations for which the neutral invasion success would just be 1/*N*.

Yet, the colonization resistance of gaseous populations still is weak compared to partially-jammed populations. Their infiltration is nearly impossible, as it is exponential in the ratio of the thickness of the jammed fraction and the thickness of the founder population, which is at most several cell diameter. The founder population at the floor of the chamber is essentially isolated from the supply channel, through the constant shedding of jammed cells acting as diffusion barrier for any invader. Invasion can only be achieved if this diffusion barrier is broken down, say by an antibiotic treatment or chamber deformation or increase in chamber flow.

Finally, we discuss the case where the growth rate of the resident is reduced by the action of an antibiotic, as done in our experiments. Invadable, gaseous, populations become invaded at rate that is increased by factor of *N_e_s*, where *s* ≪ 1 is the growth rate defect and *N_e_* ≫ 1 is the effective population size of the chamber. The situation is similar to well-mixed populations. The only difference is that, due to the spatial structure of the population, *N_e_*/*N* < 1 is mildly and strongly reduced in the gaseous and jammed states, respectively.

#### C.1 Tracking lineages backward in time

To model infiltration and neutral take over, we generalize an analysis of “gene surfing”^57^ to include a distinction between collective diffusion and self-diffusion. Suppose we sample a cell at position *ξ* at present time *τ* and seek to determine the probability density *G*(*y*, *t*|*ξ*, *τ*) that the ancestor of the cell was at position *y* at earlier time *t*. *G* then describes backward in time the position of the cell’s lineage, which is subject to self-diffusion and advection (no proliferation). *G* therefore satisfies a generalized diffusion or Fokker-Planck equation, which takes the form

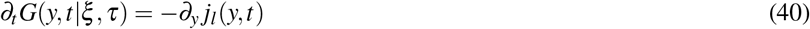

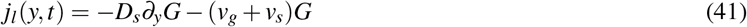

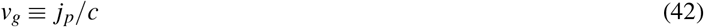

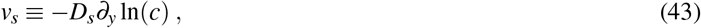

where *D_s_* is the self-diffusivity and the particle current *j_p_* is given by

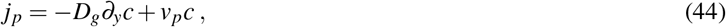

where *v_p_*(*y*, *t*) is the particle velocity at (*y*, *t*) due to any external force.

The key part here is the contribution of the lineage current due to self-diffusion. The mathematical form *v_s_* = −*D_s_∂_y_*ln(*c*) is fixed by the requirement that, for *v_g_* = 0, *G* ∝ *c* must be a steady sate solution with vanishing current.

Suppose, the chamber population has reached a steady state and is large enough so that we can ignore density fluctuations. Assuming there is no external force, *v_p_* = 0, the steady state of the ancestor distribution is then given by

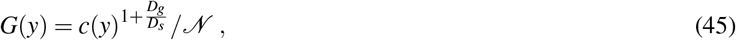

where 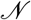 is a normalizing factor,

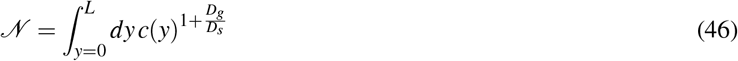

From this, we can conclude that one particle injected at *y*, will after relaxation in the cavity take up a fraction

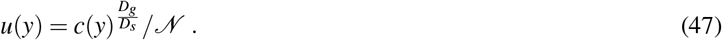

This expression is particularly useful for the gaseous phase where gradient- and self-diffusivity are nearly identical. In this case, we have an approximately cosine density profile and, with *D_g_* = *D_s_*, we obtain

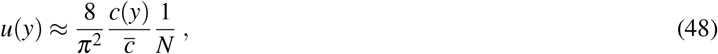

where 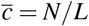 is the mean population density in the chamber. This expression shows that the neutral invasion success is much less than the well-mixed expectation 1/*N* because the invading cell has to enter from the supply where the cell density is very low.

In the case of a jammed population, the above expressions are not so useful because they require us to know the minute spatial density variations in the jammed phase. Instead, we can exploit the fact that the density hardly varies apart from a boundary layer near the exit of the chamber. A vanishing particle current at steady state requires

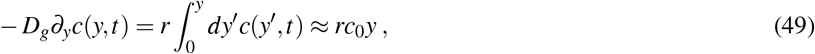

where we used *c*(*y*, *t*) ≈ *c*_0_ in the last step. Hence, the steady state ancestor distribution is a decaying Gaussian,

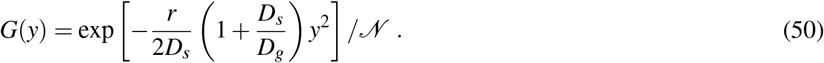

Since *D_s_*/*D_g_* ≪ 1 in the jammed phase, we conclude that fixation becomes small very rapidly when 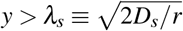.

In our experimental system, we found that *λ_s_* is about one cell diameter and the thickness of the jammed fraction was about 50 cell diameters, even just after the tipping point. It is, therefore, appropriate to think of the jammed populations as diffusively isolated from the outside environment.

## Supplementary Table

**Table 1.**
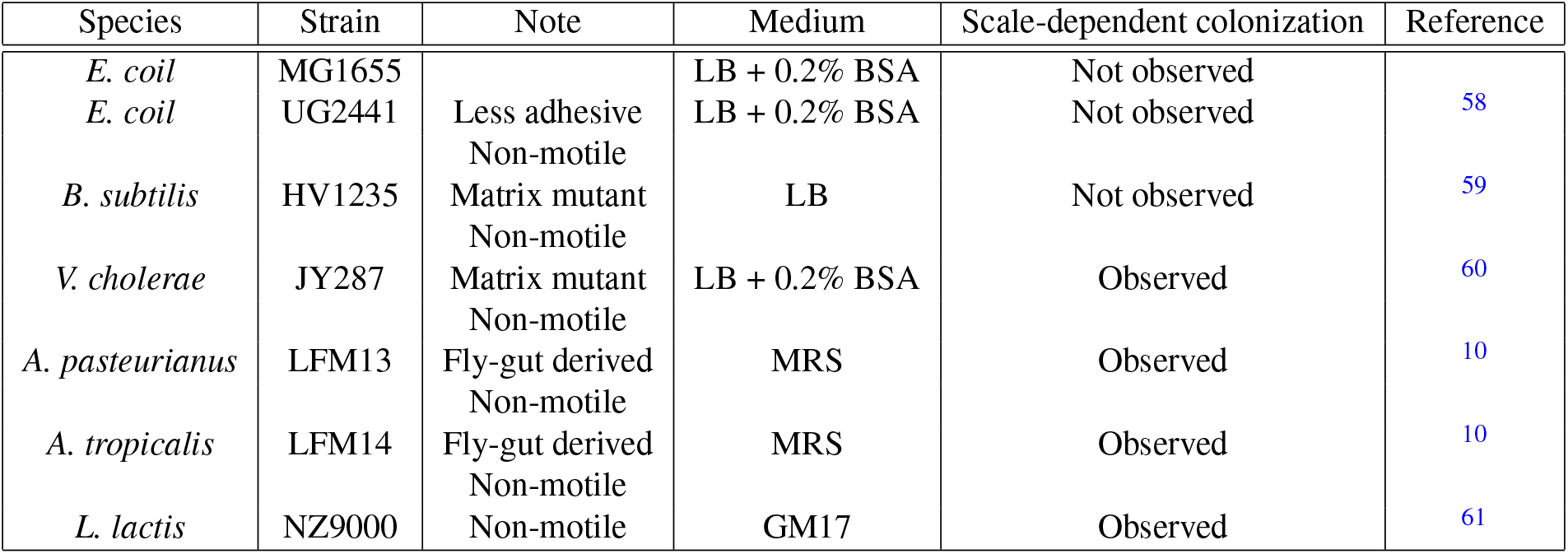
Strains cultured in panflute devices. Scale-depedent colonization was observed across multiple species (see Fig. S3), but was not observed for strains which were highly adhesive to walls or had a strong capability of bioflim formation or filamentation.

| Species | Strain | Note | Medium | Scale-dependent colonization | Reference |
| --- | --- | --- | --- | --- | --- |
| *E. coil* | MG1655 |  | LB + 0.2% BSA | Not observed |  |
| *E. coil* | UG2441 | Less adhesive<br>Non-motile | LB + 0.2% BSA | Not observed | 58 |
| *B. subtilis* | HV1235 | Matrix mutant<br>Non-motile | LB | Not observed | 59 |
| *V. cholerae* | JY287 | Matrix mutant<br>Non-motile | LB + 0.2% BSA | Observed | 60 |
| *A. pasteurianus* | LFM13 | Fly-gut derived<br>Non-motile | MRS | Observed | 10 |
| *A. tropicalis* | LFM14 | Fly-gut derived<br>Non-motile | MRS | Observed | 10 |
| *L. lactis* | NZ9000 | Non-motile | GM17 | Observed | 61 |

## Supplementary Figures

**Figure S1.**
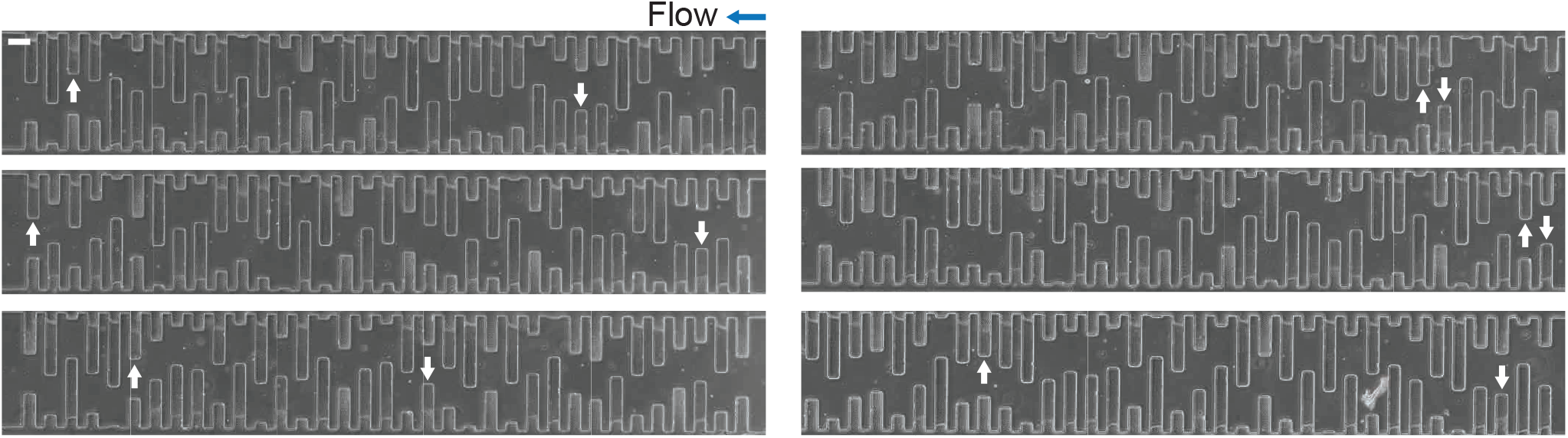
Colonization patterns in randomized panflutes. The effect of anterior populations in the same row was tested by randomizing the order of chambers. The transition to a phase-separated state was observed independent of the order of the chambers. White arrows show the onset of jamming. The scale bar indicates 100 μm.

**Figure S2.**
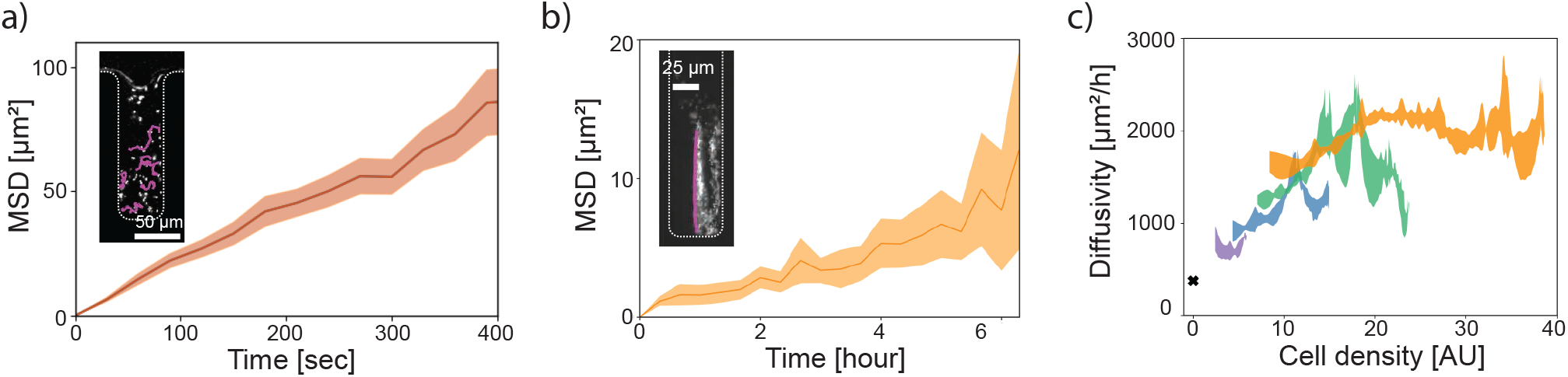
Measuring self-diffusivity and collective diffusivity. (a) Self-diffusivity was measured by tracking single cells in gaseous phases (inset: a snapshot of single-cell tracks). The self-diffusivity was calculated as 376 ±6 μm^2^/h from the mean square displacements in the horizontal direction. The error was estimated from fitting. (b) Diffusivity in jammed phases was estimated by manually tracking lineages (inset: a snapshot of a lineage). The diffusivity was calculated as 0.62 ±0.02 μm^2^/h from the mean square displacements in the horizontal direction. (c) Collective diffusivity was calculated from steady-state density profiles (see Method) of gaseous phases in 4 chambers with various depths (the colors show different chambers in the same panflute). The measured collective diffusivity showed a trend of unimodality. The black cross shows the self diffusivity measured in (a). The errors were estimated from the smoothing parameters.

**Figure S3.**
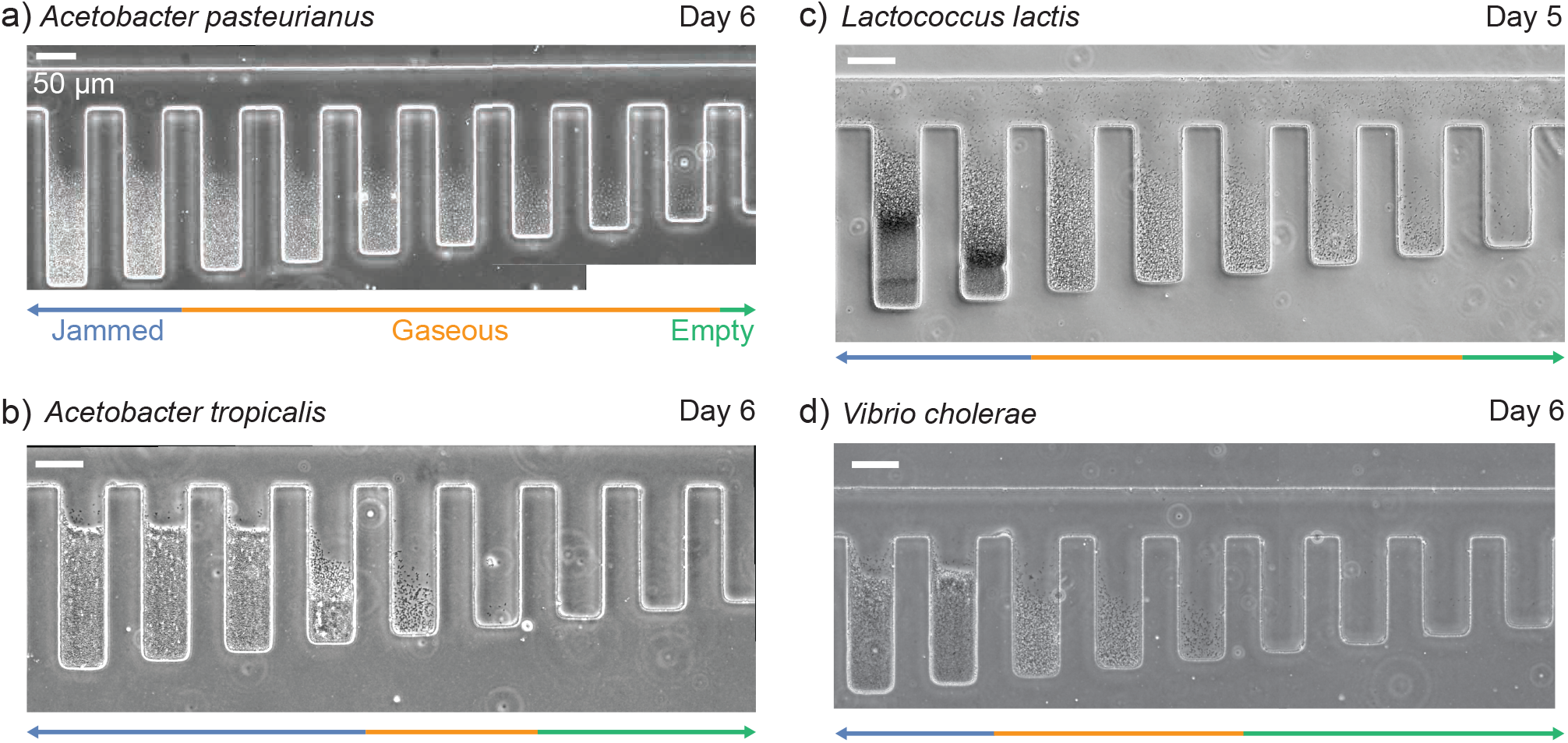
Three colonization phases are observed in different bacterial species. Pictures were taken after 5-6 days of incubation in microfluidic devices. Despite biofilm formation (b) and nutrient depletion (c), we observed qualitatively similar colonization patterns. Scale bars indicate 50 μm.

**Figure S4.**
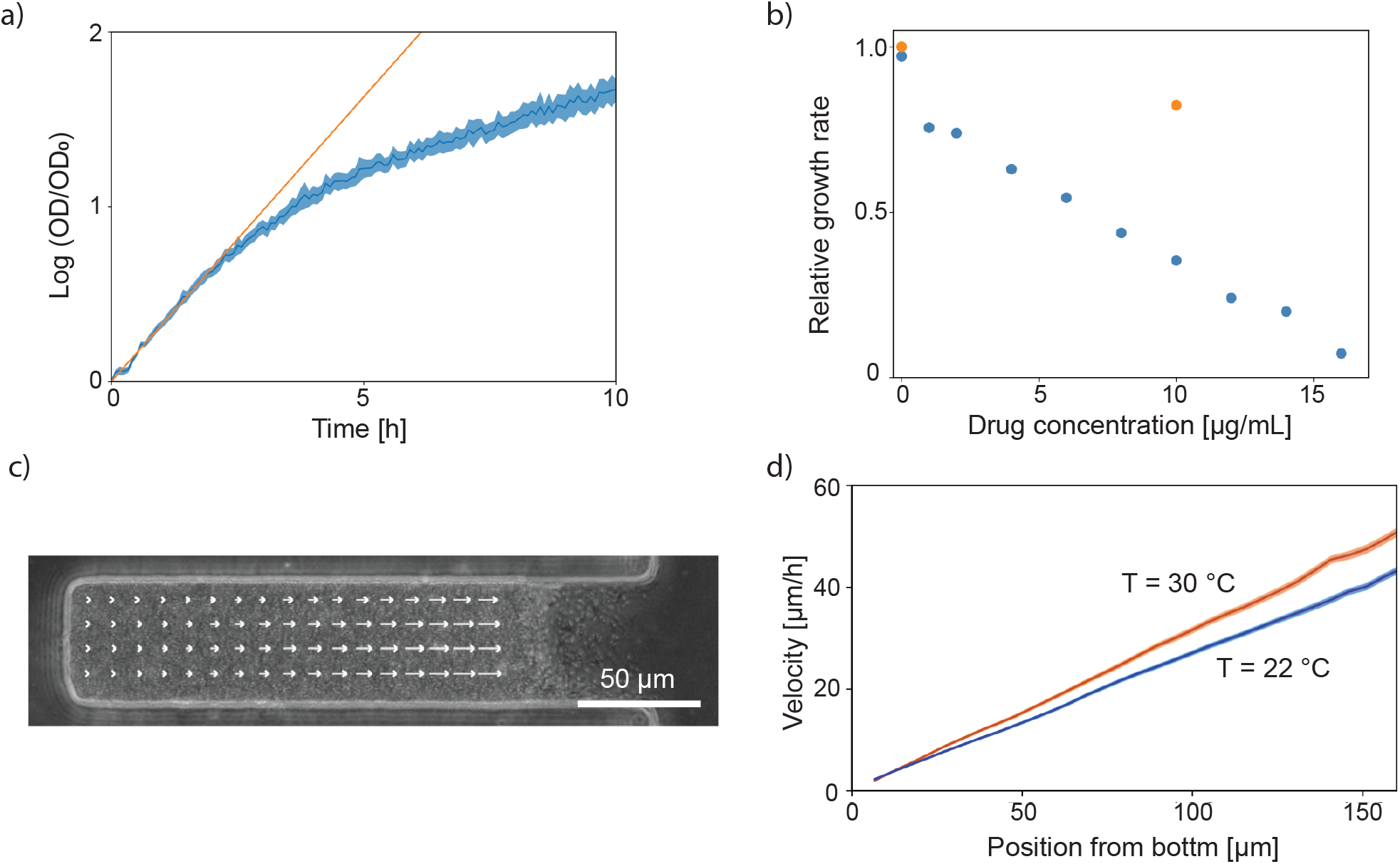
Growth rate measurements with a plate reader and particle image velocimetry (PIV). (a) The growth rate of *Acetobacter indonesiensis* was measured with a plate reader. The maximum growth rate at 30 °C was estimated as 0.325±0.003 h^−1^ from the initial growth of 4 independent populations. (b) The growth rate of tetracycline-sensitive (blue) and resistant (orange) cells was measured with various drug concentrations and normalized by the the growth rate of drug-resistant cells in the absence of the drug. The minimum inhibitory concentration was estimated about 17 μg/mL by extrapolating the plot. The averaged growth rate for each condition was calculated from 4 replicas. (c) A schematic of PIV analysis. Arrows show the local velocity of the positions. The length of arrows is proportional to the local velocity. (d) The local velocities of cells at high temperature (red, 30 °C) and low temperature (blue, 22 °C) were linear functions of the position from the bottom of a microfluidic chamber. The solid lines were the local velocities averaged over 3 hours, and the shaded regions show the standard error of mean. The growth rate of cells was derived from the slope of the linear function as 0.332 ± 0.007 and 0.280 ± 0.001 h^−1^ at high and low temperature respectively.

**Figure S5.**
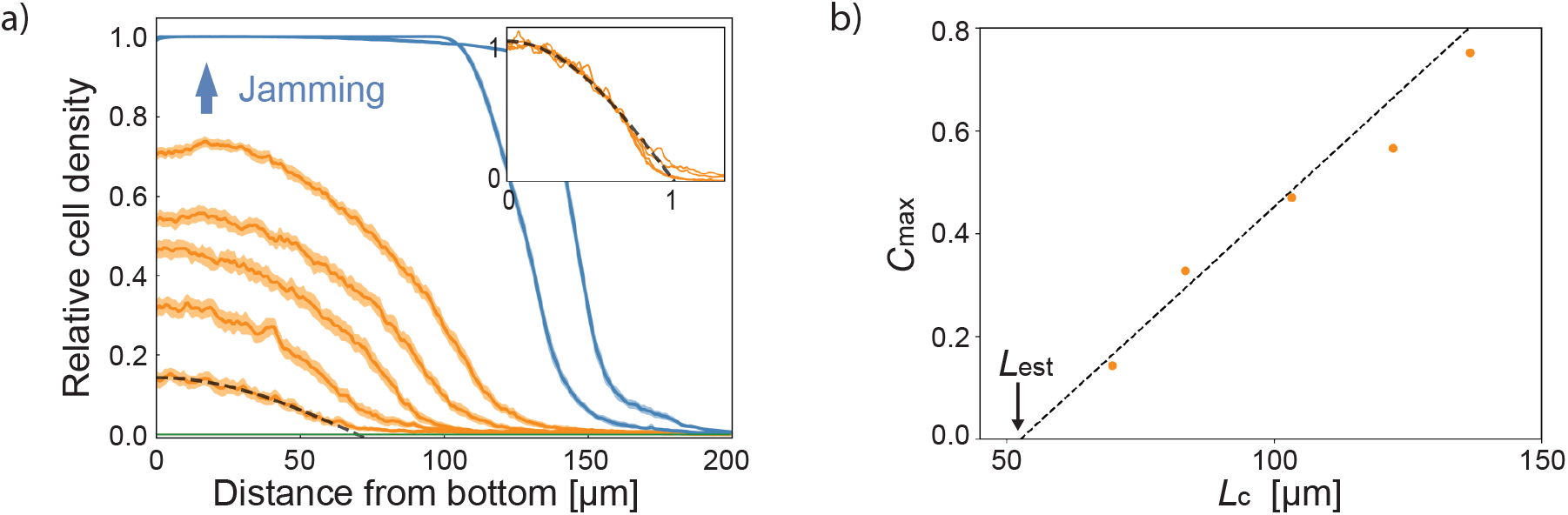
Density profiles of gaseous phases can be scaled to approximately collapse onto a master curve. (a) The steady-state density profiles in a Microfluidic Panflute device. The plot is taken from Fig. 1d. The density profiles in the gaseous state can be well approximated by the function *c*_max_ cos(*πx*/2*L_c_*), which can be seen in the rescaled plot showing *c*/*c*_max_ vs. *x*/*L_c_* (inset). (b) Plotting *L_c_* vs. *c*_max_ yielded a near linear relationship in the gaseous state. Extrapolating the linear fitting of the lowest three points to vanishing density yielded an estimate of the establishment length *L*_est_ ≈ 53 ±7 μm. The error was estimated from fitting. By comparison, our linear stability analysis predicted *L*_est_ ≈ 53 ± 1 μm (see main text).

**Figure S6.**
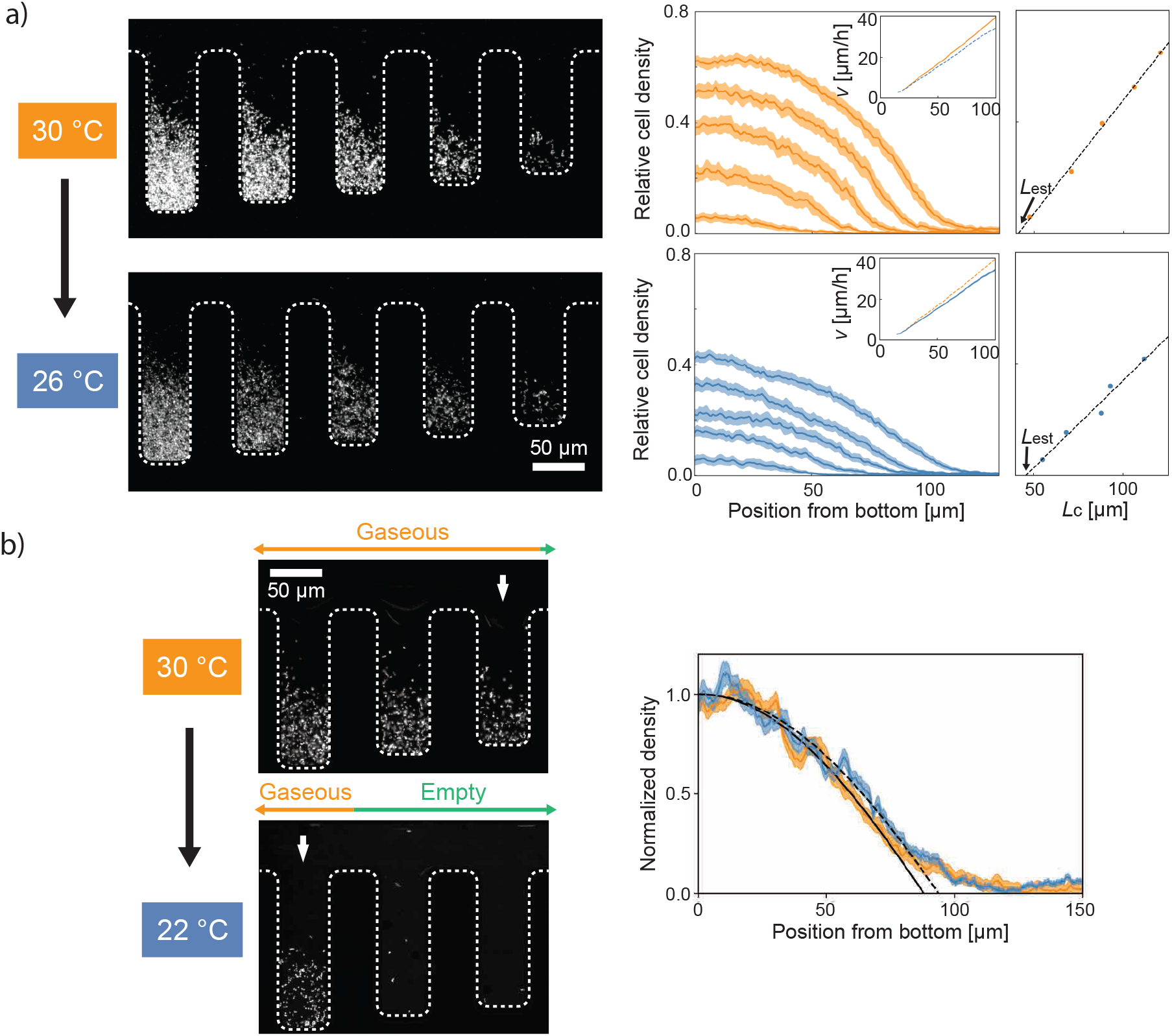
The establishment length *L*_est_ shifts upon a temperature change. We performed two temperature shift experiments, where we inoculate a panflute at one temperature and, after sufficient relaxation, shift to another temperature, after which we let the system relax again. (Relaxation often took more than 5 hrs.) (a) 30 °C to 26 °C. Right: The density profiles changed within the different crypts changed substantially the temperature change. Densities are consistently higher at 30 °C (orange) than at 26 °C (blue). The profiles were measured at steady states with fluorescent microscopy. The insets of the plots show PIV measurements whose slopes indicated the growth rates (see Fig. S4c and d). The growth rate decreased to 87.2 ± 0.8%. The shift of *L*_est_ was analyzed by determined by extrapolating the relation between cavity length and maximal population density at the floor of the crypts to vanishing cell density, similarly to Fig. S5. We found that the establishment length *L*_est_ increased by 112 ± 11 %. This change was consistent with our theory 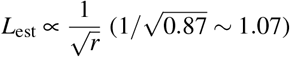. (b) The establishment length shifted upon the temperature change (from 30 °C to 22 °C). The steady-state density profiles at 30 °C (orange) and 22 °C (blue) were fitted by a cosine function (black solid and dashed lines, respectively) and normalized. The establishment length was defined by the *x*-intercept. The relative change of this critical length (6.6 %) was consistent with our theory predicting it to be given by the square root of the relative growth rate change (8.6 %, Fig. S4d). Note that, while these temperature shift experiments are consistent with a pure growth rate change, they come with the caveat that, besides growth rate, additional cell traits might be affected that influence the phase behavior, for instance, the shape of the cells or their intercellular mechanical interactions.

**Figure S7.**
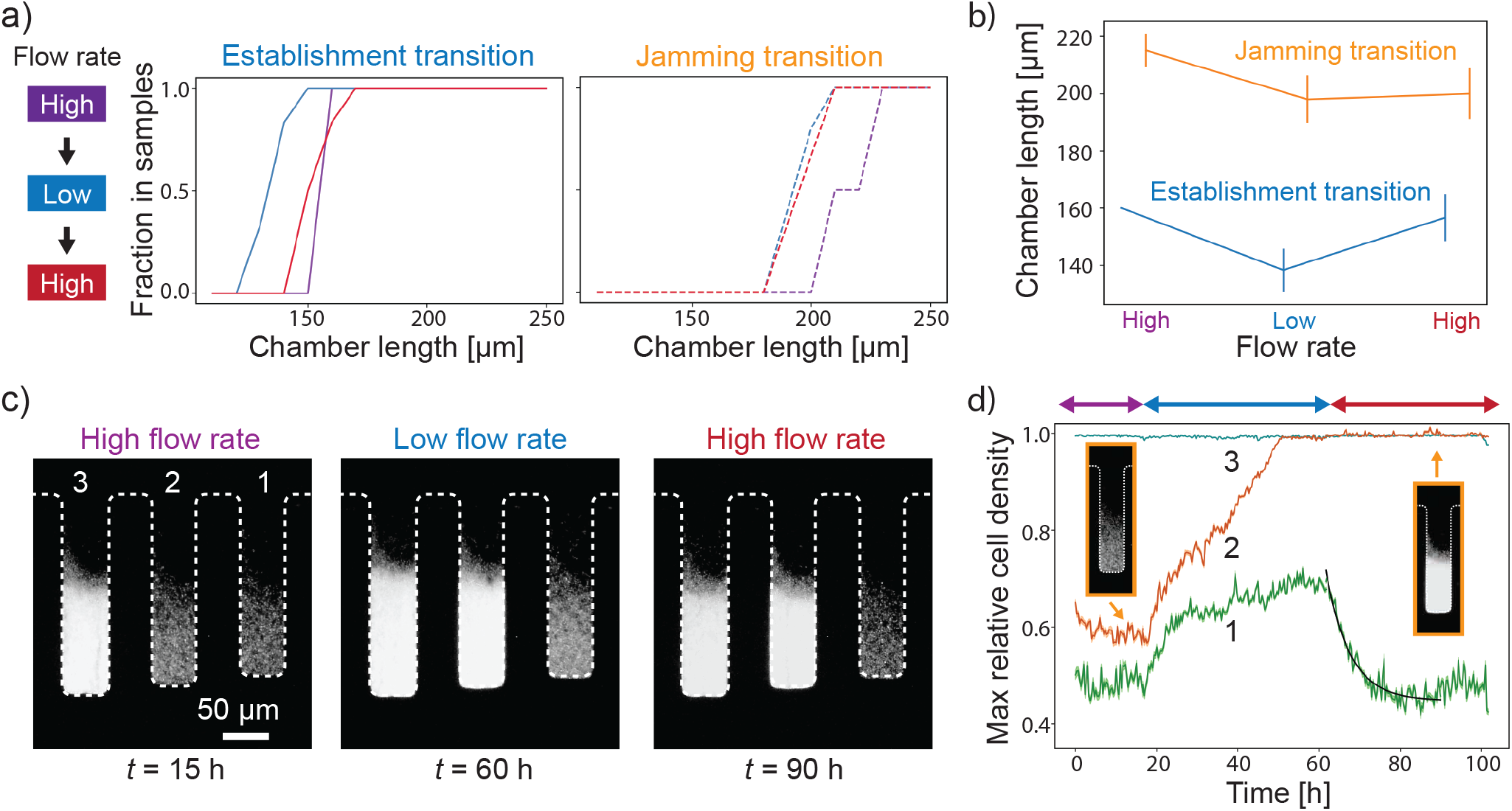
Phase shift and bistability upon flow rate change. This figure documents how colonization patterns in the Microfluidic Panflute changed as we changed flow rates from “High” (purple, 0.8 μL/h) to “Low” (blue, 0.3 μL/h) and back to “High” (red), while allowing the populations to reach steady state after each flow rate change. Note that a flow rate increase (decrease) corresponds to a decrease (increase) in the effective chamber depths (see Fig. S8). (a) The fraction of occupied chambers (left) and the fraction of jammed chambers (right) are shown as a function of chamber size (incremented by 10 μm). The lines are colored according to the state diagram (left). *n* = 3-6 for each chamber length. Note that, while the critical length for establishment (left) shifted reversibly as the flow rates was changed, we found hysteresis in the jamming transition (right). (b) The average transitional lengths extracted from (a) are displayed. The error bars show the standard error of the mean. The point without the error bar means that all samples had their establishment transition at the same (discrete) chamber length. (c,d) Time tracking of populations growing in the same Microfluidic Panflute. (c) Steady state snapshots of chambers that are near the jamming transition. Note that, while the occupancy pattern of chambers 1 and 3 changed reversibly, chamber 2 showed hysteric behavior, indicating bistability. (d) Dynamics of the maximal cell density at the floor of the chambers as the flow rate was cycled. Colored lines show the temporal dynamics of the maximum relative cell density in each chamber. The density profiles in the chambers were calculated by averaging the fluorescence across the horizontal direction at each time point. The shaded region shows the standard error of the mean. Two representative snapshots for two stable states of chamber 2 are shown in the insets. The black line shows an exponential fit to the population decay. The decay time was 5.9 ± 0.4 hour (the error was estimated from fitting).

**Figure S8.**
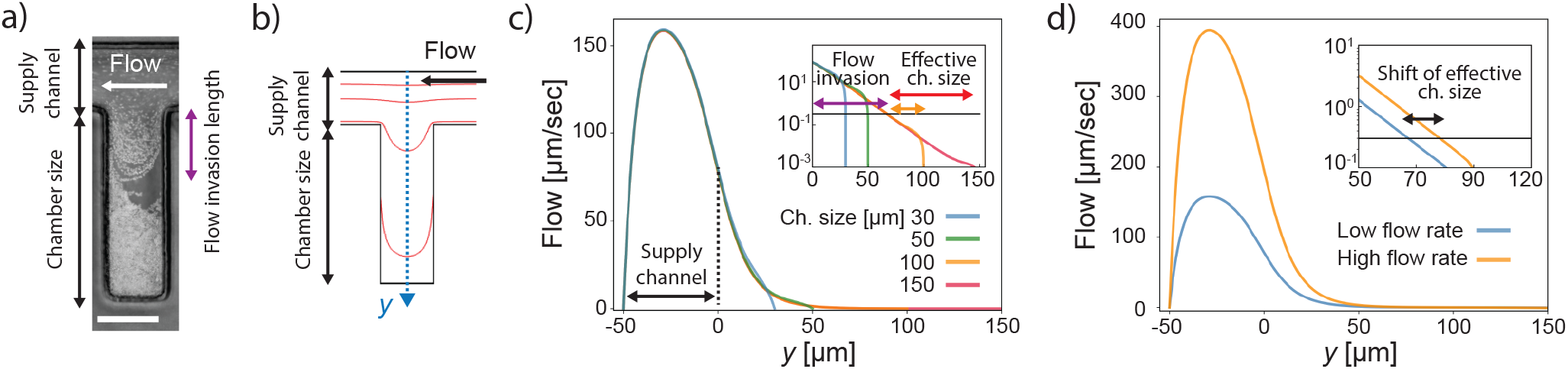
Simulations of the hydrodynamic flow fields in the Microfluidic Panflutes. (a) Streamlines of the flow were visualized by overlaying 90 frames taken every 2 seconds. The trajectory of cells showed that the typical scale of the flow invasion length was about 60 μm. The scale bar shows 50 μm. (b) The hydrodynamics in our microfluidic devices were simulated using COMSOL. Red lines show streamlines. (c) The horizontal flow velocity along the blue dotted line in (b) is shown as function of vertical position *y*. Note that the flow rapidly decays from the opening (*y* =0) towards the floor of the cavity. The inset shows the flow profiles in a semi-log scale. We define an arbitrary threshold flow velocity (0.3 μm/s, the black line in the inset) to define the flow invasion length and the effective chamber length, shown as the purple arrow and the orange (100-μm chamber) and red (150-μm chamber) arrows, respectively. The flow invasion length is constant for chamber sizes beyond 100 μm. (d) The effective chamber size gets shorter by 10 μm when the flow rate changes from low (blue, 100 μm/s average flow rate) to high (orange, 250 μm/s average flow rate), shown as the black arrow in the inset. Note that the shift of the effective chamber size is not sensitive to the choice of the threshold flow velocity.

**Figure S9.**
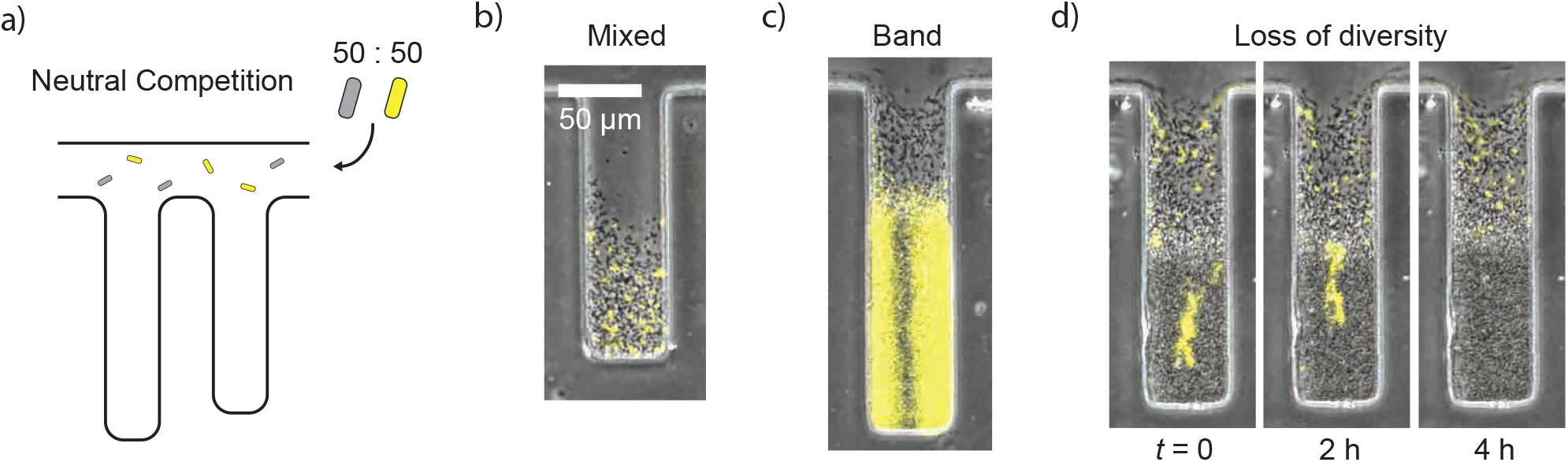
Neutral dynamics of mixed cultures reveal suppressed lineage diffusion in jammed population. (a) A schematic of neutral competition experiments. A 50:50 mixture of wild-type and labeled invader strains was inoculated into unoccupied chambers without antibiotics. (b) Labeled cells were sparsely distributed in a gaseous phase. (c) Steric interactions and proliferation produced band-like patterns in a jammed phase. The population dynamics were dominated by a small number of cells at the bottom of a cavity. (d) Diversity was rapidly lost in a jammed phase. A cluster of GFP-tagged cells was pushed out of the chamber by the population growth in a few generations.

**Figure S10.**
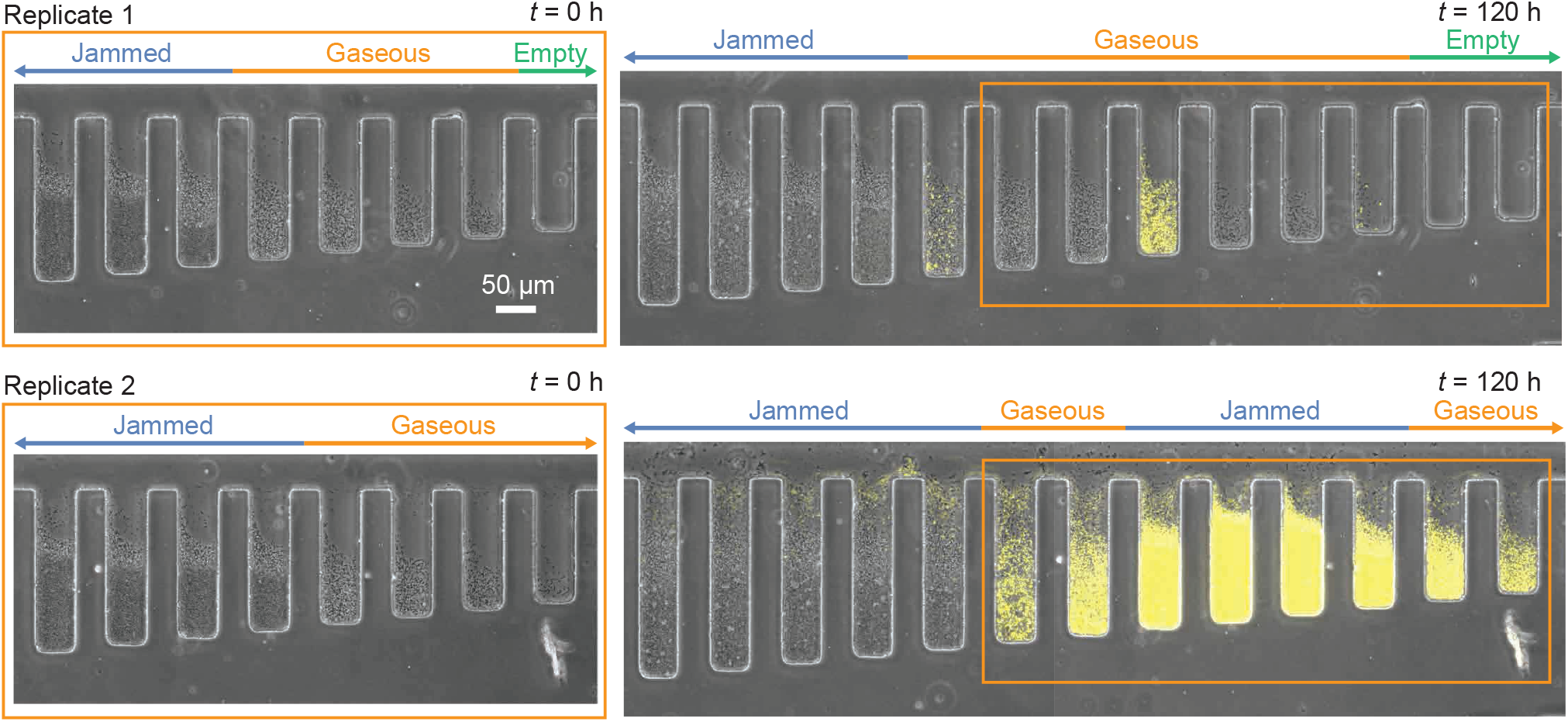
Replicas of invasion experiments with 10 μg/mL tetracycline. Replicas from other rows on the same microfluidic chip. Orange frames show the same positions. Colonization resistance of the jammed phases was consistently observed, while the rate of invasion varied across replicas (less successful in the replica 1, and more successful in the replica 2).

**Figure S11.**
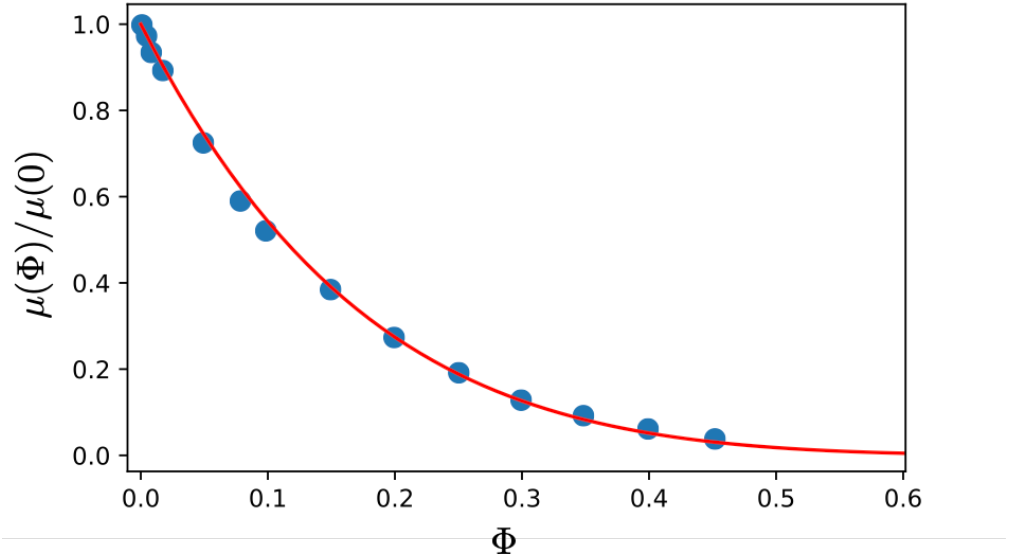
Collective mobility with complete many-body hydrodynamic interactions (blue circles) and a fit to the Richardson-Zaki scaling form (red line).

**Figure S12.**
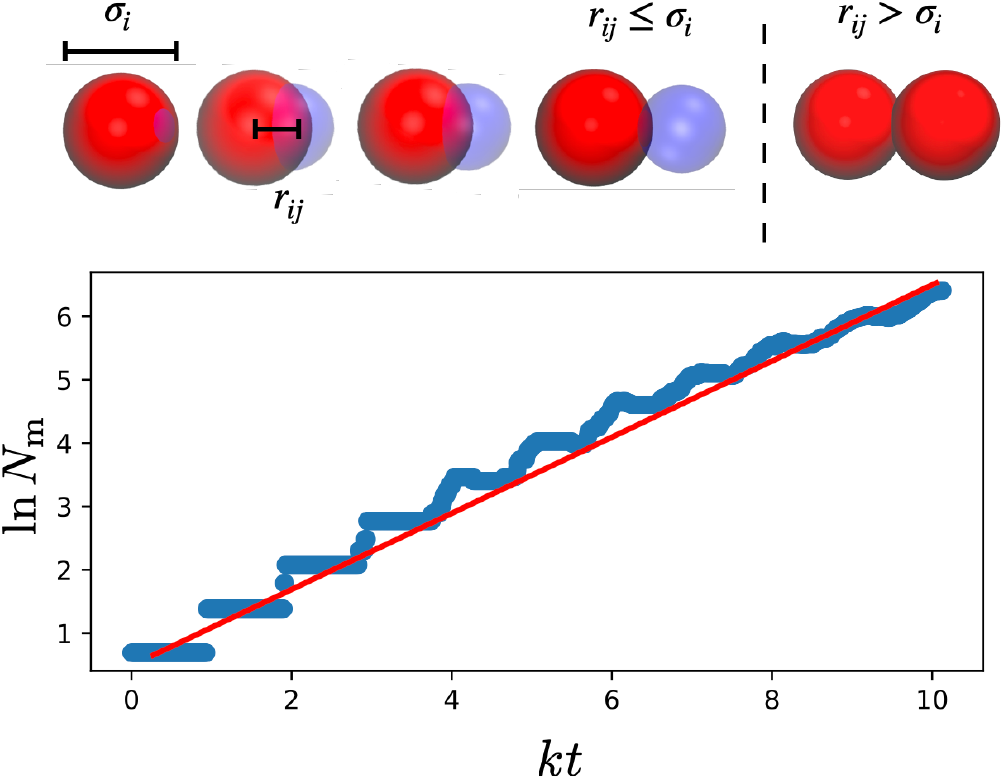
Illustration of the growth and division. (top) Illustration of the mechanical model of division of a mother (red) daughter (blue) particle pair, where the characteristic size of the mother is *σ_i_* and its displacement from a daughter is *r_ij_* (bottom) Illustration of the subsequent exponential proliferation of particles in time over 10 division times. The red line is a guide to the eye.

**Figure S13.**
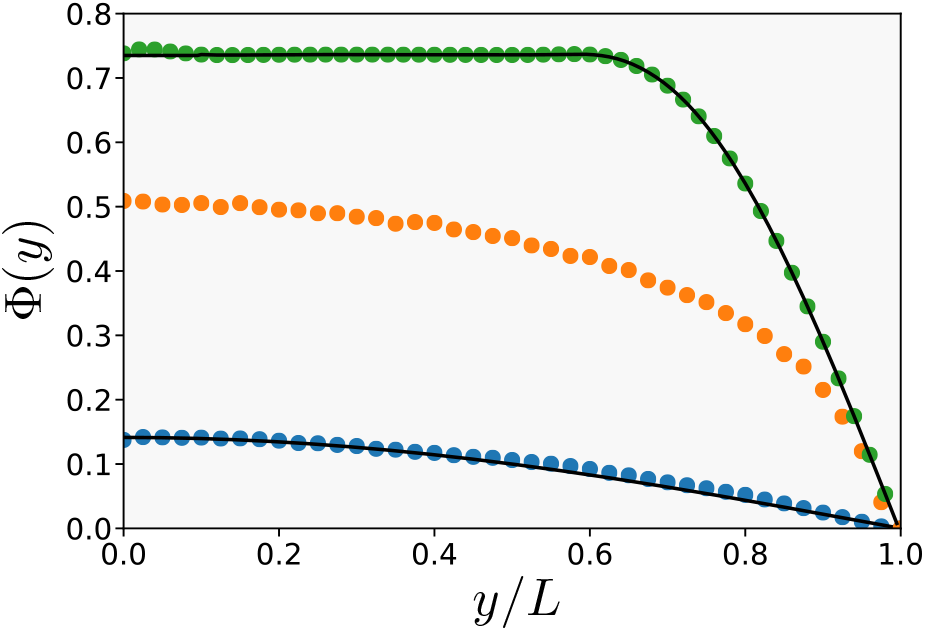
Coarse-grained packing fraction profiles computed from simulations at *L*/*L*_est_ ={1.2,2,4}(blue, orange and green) compared to the analytical predictions of the reaction diffusion model (solid lines).

**Figure S14.**
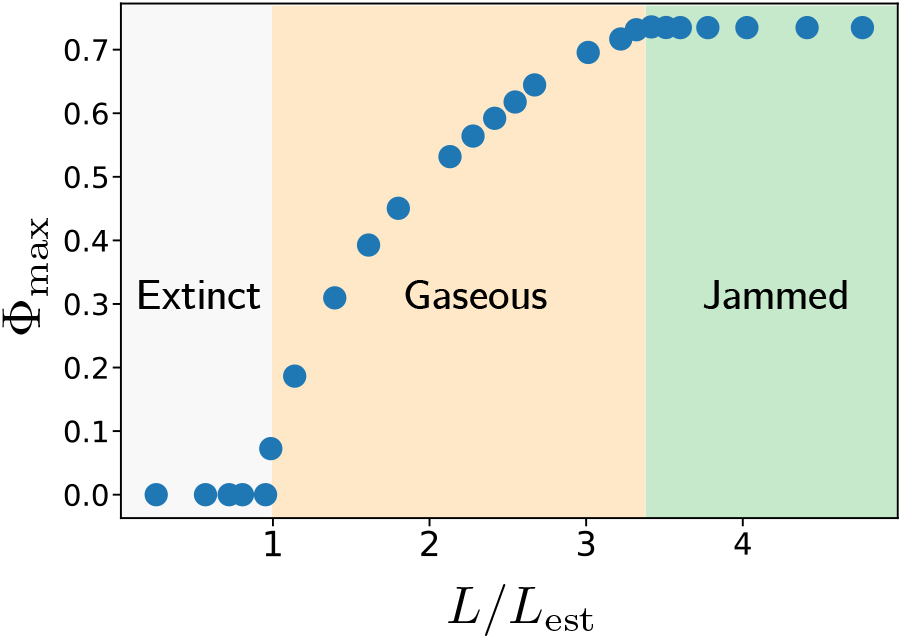
Phase diagram for the proliferating soft disks determined by the maximum coarse-grained packing fraction in the chamber.

## Footnotes

1 In our experiments, removal is dominated by outflow. Cell death can also be included through an effective growth rate, representing the difference between growth and death rate

2 The mechanism of Motility-Induced Phase Separation is based on an active movement (motility), which can generate a negative effective diffusivity^26^.

